# Schlafen-11 and -9 are innate immune sensors for intracellular single-stranded DNA

**DOI:** 10.1101/2024.02.24.581893

**Authors:** Peng Zhang, Xiaoqing Hu, Zekun Li, Qian Liu, Lele Liu, Yingying Jin, Sizhe Liu, Xiang Zhao, Jianqiao Wang, Delong Hao, Houzao Chen, Depei Liu

## Abstract

Nucleic acids are major structures detected by the innate immune system. Although intracellular single-stranded DNA (ssDNA) is accumulated during pathogen infection or pathogenesis, it remains unclear whether and how intracellular ssDNA stimulates the immune system. We report that intracellular ssDNA triggers cytokine expression and cell death in a CGT motif-dependent manner. Through a genome-wide CRISPRLCas9 screen, we identified Schlafen-11 (SLFN11), which is essential for the response to endogenous and pathogen ssDNA. SLFN11 directly binds ssDNA containing CGT motifs and translocates to the cytoplasm upon ssDNA recognition. The mice deficient in SLFN9, the homologue of SLFN11, were resistant to CGT ssDNA-induced inflammation, acute hepatitis and septic shock. This study establishes CGT ssDNA and SLFN11/9 as a novel type of immunostimulatory nucleic acids and pattern recognition receptors, respectively.

**One-sentence summaries:** single-stranded DNA inside cells can awaken Schlafens with a CGT motif to trigger cytokine expression and lytic cell death.

## Main Text

Pattern recognition receptors (PRRs) play a crucial role in innate immunity, as they recognize pathogen-associated molecular patterns (PAMPs) and damage-associated molecular patterns (DAMPs) and initiate innate immune responses, such as type I interferons, inflammation and cell death (*1, 2*). The first class of PRRs to be identified was the Toll-like receptors (TLRs), which are membrane proteins that recognize extracellular or endosomal pathogen ligands. However, PAMPs and DAMPs can also exist in cytoplasm or nucleus (*1–3*). Consequently, most TLRs have intracellular complementary PRRs that sense similar ligands, such as TLR3 and RIG-I/MDA5 for double-stranded RNA (*1–3*). TLR9 is a specific receptor for CpG motifs in single-stranded DNA (ssDNA) that is internalized into endosomes, but it is ‘blind’ to cytosolic or nucleic ssDNA and is primarily expressed in B cells and plasmacytoid dendritic cells (*1, 2, 4*), suggesting the existence of a broadly expressed sensor for intracellular ssDNA (*5*). However, such a general intracellular ssDNA sensor that initiates innate immune responses has not been unequivocally identified.

Emerging evidence indicates that intracellular ssDNA accumulates due to pathogen infection (*6–8*), Trex1 mutation (*9*), replication stress (*10*), chromosome instability (*11*), dysregulation of retrotransposon (*12*), R-loop accumulation (*13*), and gene therapy (*14*). Although the dsDNA-specific sensors cGAS (*15, 16*) and IFI16 (*17*) have been reported to mediate cytosolic ssDNA-induced type I interferon responses, they actually recognize double-stranded complementary stretches of ssDNA, rather than single-stranded structure (*16, 18, 19*). Therefore, whether intracellular ssDNA can stimulate innate immune responses and contribute to pathogenesis remains obscure and controversial (*3, 8, 13, 17, 20–22*).

### Intracellular ssDNA stimulates immune responses in a sequence-specific manner

Most previous studies examining the immunostimulatory activities of intracellular ssDNA have relied on transfecting one or a few synthetic ODNs (referred to hereafter as single-stranded oligodeoxynucleotides) (*8, 17, 20–22*), potentially overlooking sequence-specific effects. To ensure comprehensive sequence coverage, we used heat-denatured *Escherichia coli* (*E. coli*) genomic DNA (gDNA) (*23*) as representative nonself ssDNA. dsDNA-preferred ethidium bromide staining and SYBR Green melting curve analysis revealed that there was a high proportion of ssDNA in the heat-denatured gDNA (*18*) (fig. S1A and B). To eliminate potential interference from TLR9 and cGAS, we transfected gDNA into HEK293 cells lacking these pathways (*24*) (fig. S1C). We observed that bacterial ssDNA decreased cell viability and increased expression of *TNF* and *CXCL8* to a greater extent than dsDNA (fig. S1, D to F), suggesting immunostimulatory properties of intracellular ssDNA. Hydroxyurea (HU) treatment-induced accumulation of endogenous ssDNA has been reported to activate cytokine expression via the cGAS-STING pathway (*10*). However, in cGAS-STING pathway-deficient HEK293 cells, we found that HU treatment still stimulated cytokine expression in a manner dependent on the accumulation of endogenous ssDNA (Fig. 1, A to C and fig. S1, G to I), as evidenced by the attenuated response when Trex1, a ssDNA exonuclease (*9*), was overexpressed (*13*). These results suggest that both transfected ssDNA and endogenous ssDNA can trigger immune responses independently of TLR9 or cGAS.

**Fig. 1.**
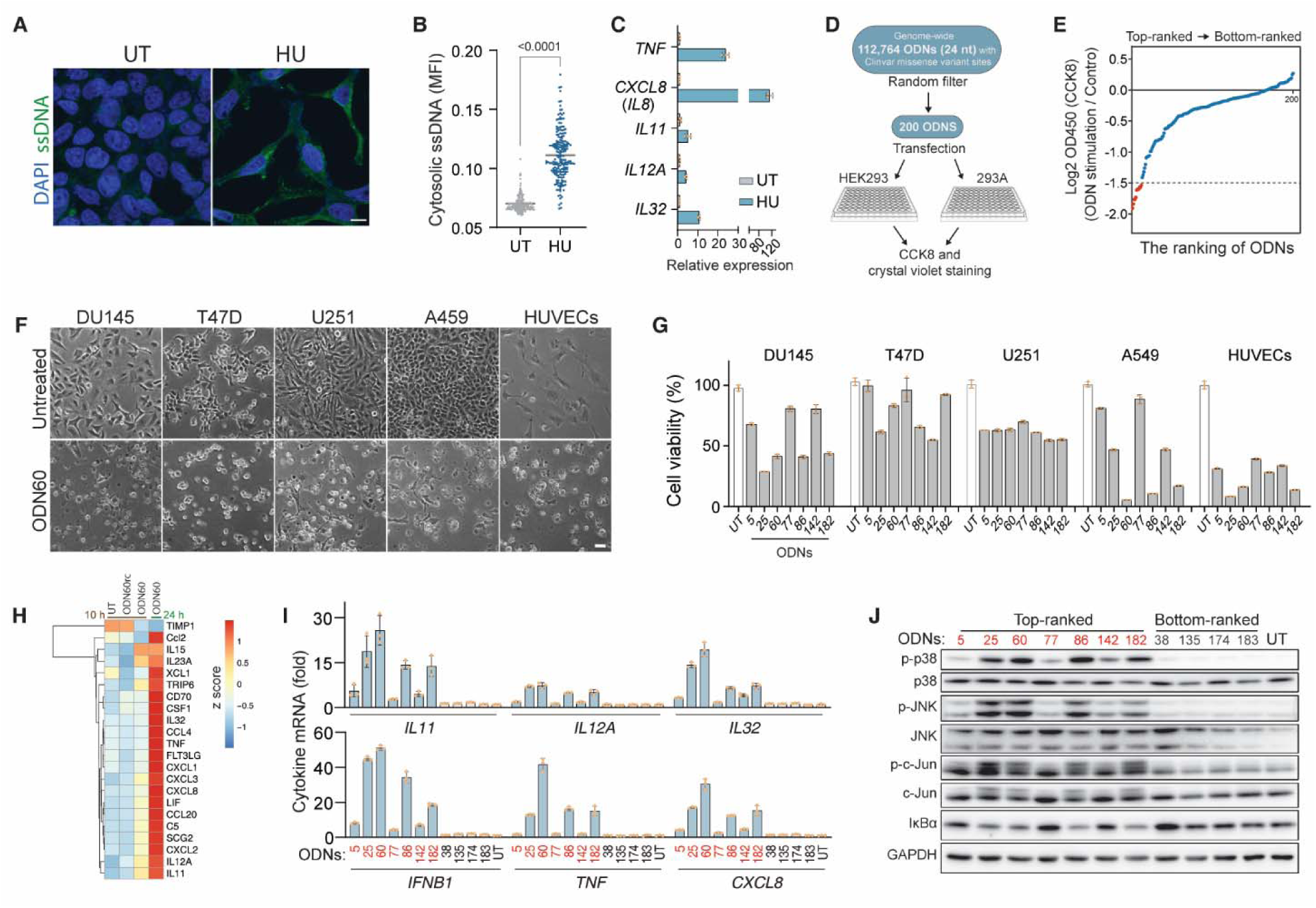
Intracellular minimal ssDNA screening reveals sequence specificity for its immunostimulatory activity. (**A** and **B**) Immunofluorescence detection of ssDNA in untreated (UT) and 4 mM HU-treated HEK293 cells (A) and mean fluorescence intensity (MFI) per cells (B). (**C**) Quantitative PCR (qPCR) analyses of indicated cytokine mRNAs in HEK293 cells exposed to 4 mM HU. (**D** and **E**) Diagram illustrating the screening of an ODN library for immunostimulatory activity (D). A total of 112,764 ODNs (24 nt) were generated based on missense SNP sites in the ClinVar database (GRCh37/hg19). Out of these, 200 ODNs (180 nM) were randomly selected and transfected into HEK293 and 293A cells (Data S1). ODNs were ranked according to the average fold change in CCK-8 assay-based cell viability of HEK293 and 293A cells (E). Top-ranked ODNs that decreased cell viability are highlighted in red. (**F** and **G**) The immunostimulatory effects of the top-ranked ODNs on the indicated cell types were assessed by morphological examination under microscopy (F) and ATP-based cell viability (G). (**H**) Heatmap representing cytokine expression profiles in 293A cells subjected to the indicated treatments. (**I** and **J**) DU145 cells were transfected with the indicated ODNs for 24 hours. The indicated cytokine mRNA (I) and immune-related signal pathways (J) were examined by qPCR and immunoblotting, respectively. Scale bar, 10 μm (A) and 50 μm (F). Data are shown as mean ± s.d. [(B), (C), (G) and (I)] from three technical replicates, and are representative of two [(E) to (G)] or there [(A to C), (I), and (J)] independent experiments. Statistical significance was determined by unpaired two-tailed Student’s t test (B).

To screen for minimal ssDNA mimics that activate immune responses, we prepared a genome-wide 24-nucleotide (nt) ODN library (Fig.1D). This ODN length was chosen because endogenous ssDNA in humans mainly ranges from 20 to 120 nt in length (*25*) and 24-nt ODNs are too short to activate dsDNA sensors through potential base-pairing stretches (*3*). Using an optimized ODN transfection method (fig. S2A), we transfected the library into HEK293 and 293A cells and observed that 14 out of 200 ODNs decreased the average cell viability of these cells to less than 40% (Fig.1E and fig. S2, B and C). We then validated these top-ranked (active-high) ODNs in a concentration gradient and found that ODN60 exhibited the strongest immunostimulatory potency (fig. S2D). Therefore, ODN60, as well as the other top-ranked ODNs, was selected as a representative stimulatory ssDNA mimic for subsequent studies. Notably, the immunostimulatory potency of ODN60 was evident at as low as 24 nM, while its reverse complementary ODN (ODN60rc), as a control, did not affect cell viability, even at 120 nM (fig. S2E). These results suggest that the immunostimulatory activities of intracellular ssDNA are highly dependent on specific sequences.

To assess whether our findings in HEK293 and 293A cells are applicable to other cell types, we transfected the top-ranked ODNs into various types of human cells. We observed that ODN-responsive cells included DU145, T47D, U251, and A549 cells and HUVECs (Fig. 1, f and G and fig. S2F), but not HeLa or 293T cells (data not shown). The stimulatory activity was exclusive to ODNs and was not observed with oligoribonucleotides (ORNs) or double-stranded ODNs (dsODNs), and was not restricted to a length of 24 nt (fig. S2, G to I). Both electroporation and transfection, but not incubation, rendered the top-ranked ODNs active (fig. S2, J and K), suggesting that the intracellular presence of ssDNA is necessary for their immunostimulatory activities. Finally, the observed morphological features (Fig. 1F and fig. 2, F and G) and live cell imaging (movie S1) revealed that the decrease in cell viability induced by stimulatory ssDNA resulted from extensive lytic cell death, which was further confirmed by annexin V and propidium iodide (PI) staining assay (fig. S2L).

**Fig. 2.**
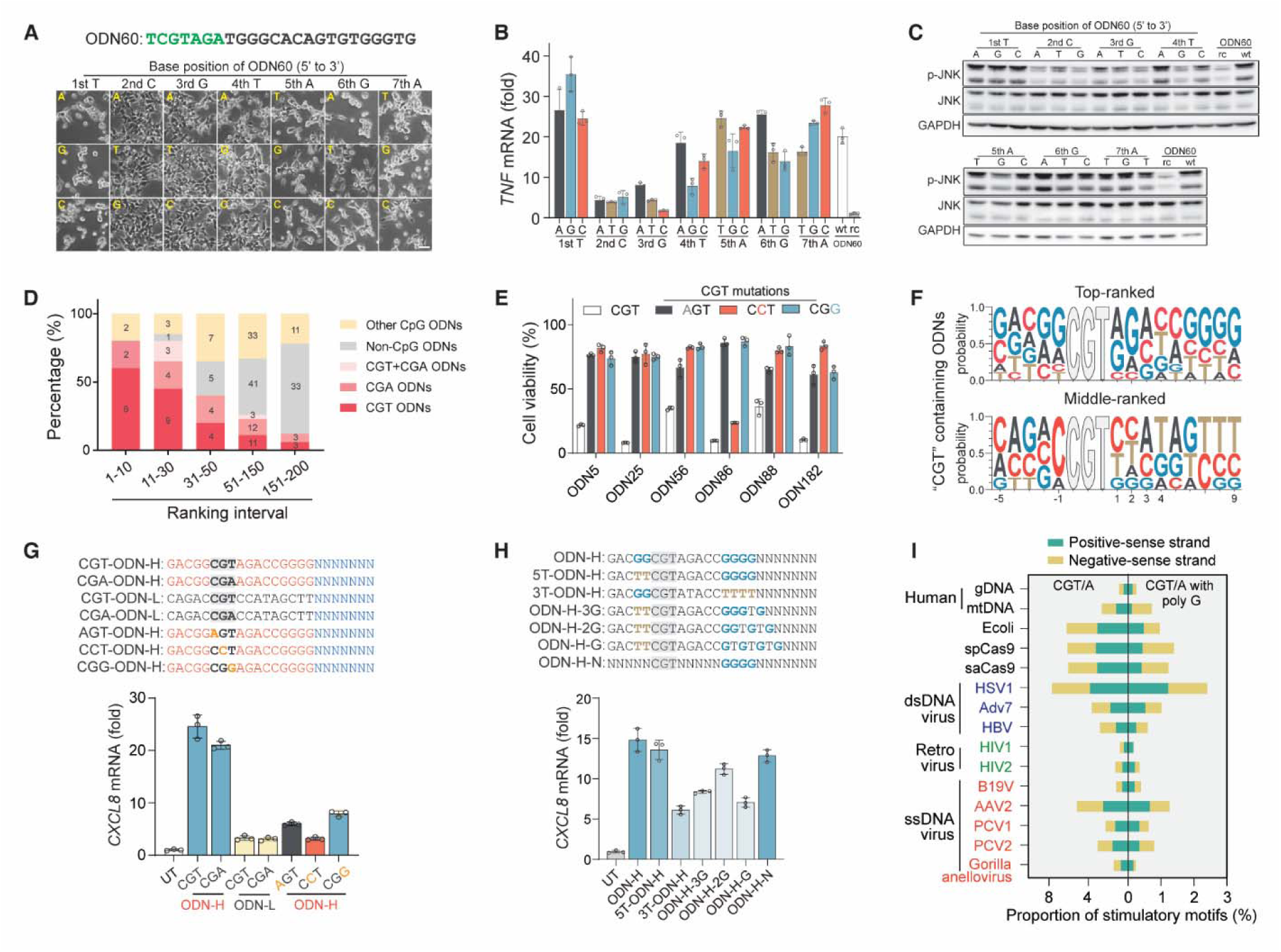
CGT/A motifs followed by a poly G are the immunostimulatory sequences for intracellular ssDNA. (**A** to **C**) HEK293 cells were transfected with ODN60 (120 nM) bearing the indicated single nucleotide mutation. Cell morphology (A) was examined 36 hours post-transfection, and *TNF* mRNA levels (24 hours) and JNK and p-JNK protein levels (12 hours) were analysed by qPCR (B) and immunoblotting (C), respectively. (**D**) Percentage of ODNs containing CGT/A motifs in the indicated ranking intervals according to Fig. 1B. (**E**) The ATP-based cell viability of HEK293 cells transfected with the top-ranked ODNs harbouring the indicated mutations in the CGT/A motif. (**F**) CGT motif-flanking sequence features (5 nt upstream to 9 nt downstream) from the top- and middle-ranked CGT ODNs were analysed with the WebLogo application. (**G** and **H**) ODN-H and ODN-L were synthesized based on the top- and middle-ranked sequence features in (F), respectively. *CXCL8* mRNA levels were measured in HEK293 cells stimulated by ODN-L, ODN-H, or ODN-H with the indicated mutations in CGT motif (G) or in the CGT flanking sequence (H) at 120 nM for 24 hours. (**I**) The proportion of CGT/A motifs, and the subset of these motifs followed by poly G sequence, in the indicated types of gDNA. The results of the cell viability assay (E) and qPCR [(B), (G), and (H)] are shown as the mean ± s.d. from three technical replicates. The data shown are representative of three [(A to C), (E), (G), and (H)] independent experiments.

### The cytokine activation further defines sequence-specific immunostimulatory capacity of intracellular ssDNA

Given that cell death is not a specific indicator for assessing immune responses, to further define the immunostimulatory activities of intracellular ssDNA, we conducted RNA-seq assays with 293A cells after 10 and 24 hours of stimulation with ODN60 or ODN60rc. As expected, only ODN60, but not ODN60rc, triggered the expression of immune-related genes compared to their expression in the untreated control (fig. S3, A and B). Importantly, the expression of many cytokines and chemokines (hereafter ‘cytokines’) was stimulated by ODN60 (Fig. 1H), which further revealed the immunostimulatory properties of intracellular ssDNA, as the transcriptional activation of cytokines is a major component of PAMP- or DAMP-induced innate immune responses (*1, 2*). Consistently, the cytokines that were triggered by endogenous ssDNA were also stimulated by the top-ranked ODNs (Fig. 1, C and I), further supporting the notion that these ODNs act as ssDNA mimics. Notably, *IFNB1*, which could not be triggered in HEK293 and 293A cells, was nonetheless stimulated in DU145 cells. Similar to the induction of cell death (fig. S2K), cytokine expression was only stimulated by intracellularly delivered ODNs, rather than incubated ODNs (fig. S3D).

Moreover, in addition to the innate immune responses signified by the activation of cytokines, we found that most top-enriched signaling pathways in ODN60-treated cells were related to immune responses (fig. S3C). In line with this, ODN60 robustly activated the JNK, p38, and nuclear factor-κB (NF-κB) signaling pathways (fig. S3, E and F), which are also typically activated by extracellular CpG ODN (*26*). Furthermore, the top-ranked ODNs also activated the JNK, p38, and NF-κB pathways in DU145 cells (Fig. 1J). To further test the activation of the NF-κB pathway, we examined the nuclear translocation of all NF-κB subunits and observed that p65 and NF-κB1 translocated into the nucleus upon stimulation with the top-ranked ODNs but not the bottom-ranked ODNs (fig. S3G). Knockdown of p65 and NF-κB1 diminished the activation of *IFNB1* and *CXCL8* induced by ODN60 (fig. S3H), suggesting that the NF-κB pathway is involved in cytokine activation induced by intracellular ssDNA. Taken together, the activation of cytokine and immune pathways further revealed the sequence-specific immunostimulatory activities of intracellular ssDNA.

### The immunostimulatory activity of intracellular ssDNA is determined by a CGT motif

Before investigating the potential immunostimulatory sequence patterns in the top-ranked ODNs, we first ruled out the possibility that differences in immunoactivity between the top-ranked and bottom-ranked ODNs were due to varying transfection efficiencies (fig. S4A). Subsequently, we designed a series of ODN60 mutations and truncation constructs. Switching the positions of adjacent bases from three pairs in the 5′ terminal region, rather than in the 3′ terminal region, led to inactivation of ODN60 (fig. S4B). In alignment with this, 3′ terminal truncations of approximately 8 nt had little impact on ODN60 immunoactivity (fig. S4D). Notably, switching only the first pair of adjacent nucleotides was sufficient to dramatically reduce the immunoactivity of ODN60 (fig. S4C). These results suggest that the immunoactivity of ODN60 was predominantly determined by its 5′ terminal sequence and sensitive to single-nucleotide changes.

To further identify the stimulatory motifs, we systematically substituted the 1^st^ to 7^th^ nt at the 5′ terminus of ODN60 with the other three types of nucleotides and evaluated the resulting activities by assessing cell viability, activation of the *TNF* cytokine and JNK pathway (Fig. 2, A to C). These results consistently showed that the 2^nd^ and 3^rd^ nucleotides were highly restricted to C and G, respectively, and the 4^th^ nucleotide, T, displayed some tolerance to A, indicating that CpG motifs were unexpectedly required for not only extracellular ssDNA (*23*), but also intracellular ssDNA to activate immune responses. We then analysed the presence of CpG motifs in the 200 screened ODNs (Fig. 1, D and E) and found that the preferred motifs are CGT and CGA (CGT/A) rather than other types of CpG motifs in the top-ranked ODNs (Fig. 2D). In line with this analysis, any single-nucleotide mutation in the CGT/A motifs led to the loss of the immunostimulatory activity of the top-ranked ODNs (Fig. 2E and fig. S4, E and F). Hence, the presence of CGT/A motifs is essential for highly immunostimulatory ODNs.

Since not all ODNs containing CGT/A motifs were potent enough (Fig. 2D), we hypothesized that the flanking sequence of CGT/A motif may also influence the potency of ODNs. To identify the pattern of flanking sequences of the CGT/A motifs with high stimulatory potential, we analysed the base composition of all CGT ODNs and aligned the flanking sequences from top-ranked and middle-ranked CGT ODNs, respectively. As most bottom-ranked CGT/A ODNs lacked downstream flanking sequences (fig. S4G), they were excluded from this analysis. We observed that the flanking sequences in the top-ranked CGT ODNs were G-rich, whereas those in the other CGT ODNs were T-rich (Fig. 2F and fig. S4H). We then synthesized ODN-H and ODN-L with a CGT/A motif encompassed by the top-ranked and middle-ranked flanking sequences, respectively. As expected, only ODN-H triggered *CXCL8* expression, while ODN-L did not (Fig. 2G). Furthermore, substitutions of G with T and the disruption of G consecution downstream of the CGT/A motif attenuated the immunostimulatory activity of ODN-H (Fig. 2H), suggesting that the CGT/A downstream poly G is important for highly active CGT/A motifs. This also explains why CGT/A motifs in the bottom-ranked ODNs are often found within the last 10 nt of the 3′ terminus, leaving insufficient room for downstream poly G tracts (fig. S4G).

To test the applicability of this motif rule for predicting the immunostimulatory potency of single-stranded gDNA, we analysed the prevalence of the identified stimulatory motifs in various types of gDNA (Fig. 2I), including human gDNA, human mitochondrial DNA (mtDNA), *E.coli* gDNA, herpes simplex virus 1 (HSV1), adenovirus type 7 (Adv7), hepatitis B virus (HBV), human immunodeficiency virus 1 and 2 (HIV1 and HIV2), human parvovirus B19 (B19V), adeno-associated virus 2 (AAV2), porcine circovirus type 1 and 2 (PCV1 and PCV2), and gorilla anellovirus. We found that *E.coli* gDNA is abundant in the stimulatory motifs, which accounts for the heightened stimulatory activity of *E.coli* genomic ssDNA (fig.S1, D to F). Notably, despite being human codon-optimized, Cas9 orthologues still possess a high proportion of the stimulatory motifs, which provides a possible explanation for innate immune activation in rAAV gene therapy (*14*). Interestingly, in comparison to dsDNA viruses, most retroviruses and ssDNA viruses that we examined possess fewer of these stimulatory motifs. However, AAV2 is notable for its high abundance of such motifs in its genome, indicating a strong stimulatory potential of its gDNA. When we introduced various types of gDNA into 293A cells via electroporation, their stimulatory potency followed the prevalence of stimulatory motifs in the gDNA (Fig. 3I).

**Fig. 3.**
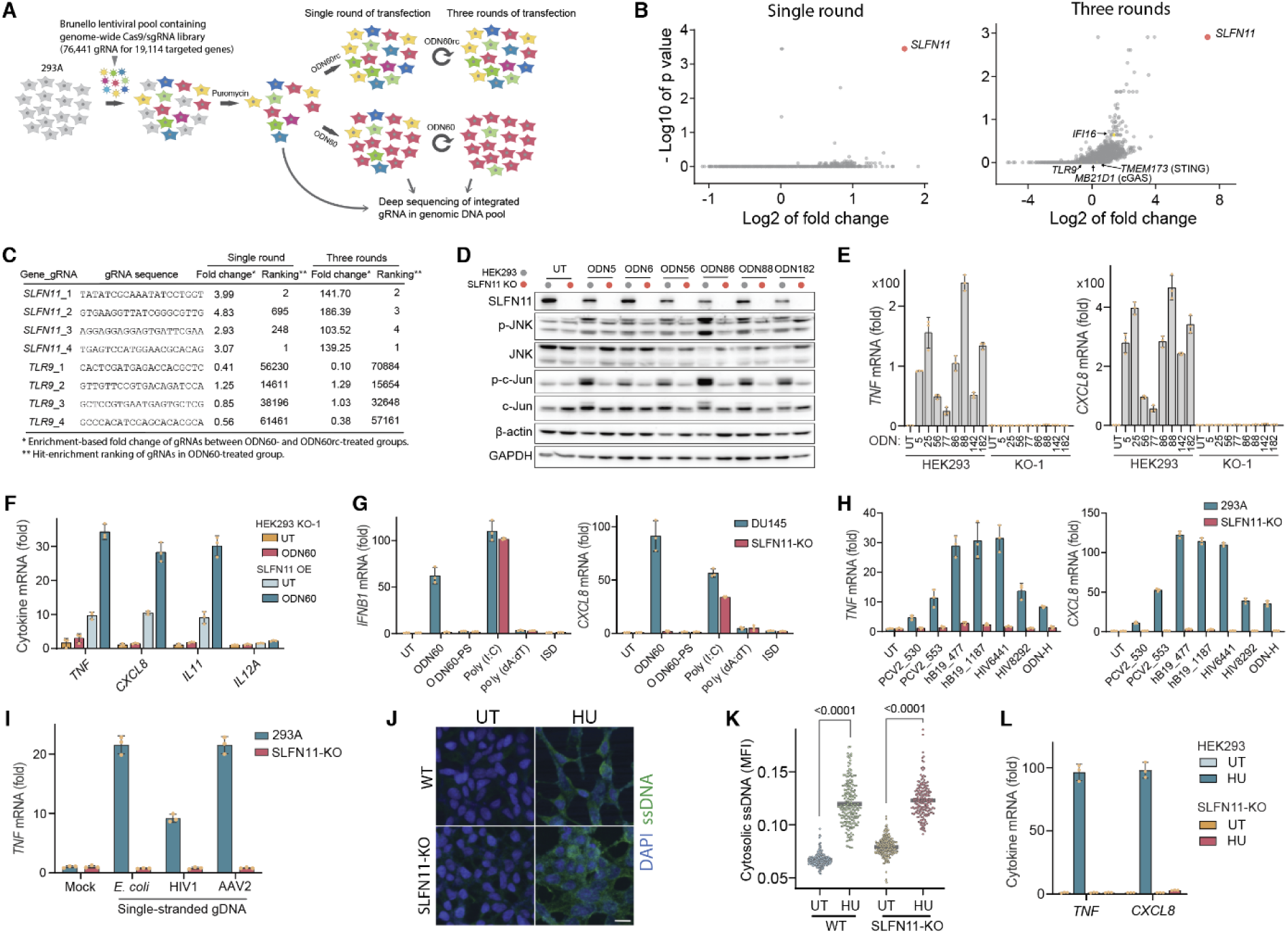
Genetic screens identify SLFN11 as an essential mediator of intracellular ssDNA-induced innate immune responses. (**A** to **C**) Schematic overview of genome-wide CRISPR screening for essential genes in response to ODN60 (A). HEK293 cells were infected with a Brunello lentiviral pool containing Cas9 and a gRNA library. After selection with puromycin, infected cells were divided into an untreated control group, an ODN60rc-transfected group and an ODN60-transfected group, and cultured for 48 hours. For three rounds of transfection screening, the cells underwent transfection for 36 hours per round, with 36 hours rest intervals between rounds of transfection. The abundance of gRNAs for each gene in living cells was determined by deep sequencing. Screen results for the enrichment of genes (B) and the gRNAs targeting *SLFN11* or *TLR9* (C) are shown. Detail data can be found in data S2. (**D**) The effects of *SLFN11* knockout on activation of the indicated pathway by CGT ODNs were analysed by immunoblotting. (**E** to **I**) Effects of *SLFN11* knockout (E to H) or rescue expression (F) on cytokine expression induced by CGT ODNs (E and F), other types of nucleic acids (G), viral genome sequence-derived CGT ODNs (H) and the gDNA from *E.coli*, HIV1, and AAV2 (I). The indicated types of nucleic acids were transfected (E to H) or electroporated (I) into cells at a concentration of 1 μg/ml. (**J** to **L**), Immunofluorescence detection of ssDNA in WT and *SLFN11*^-/-^ HEK293 cells exposed to 4 mM HU (J), scale bar, 20 μm; mean fluorescence intensity of intracellular ssDNA per cells (K) and the mRNA level of *TNF* and *CXCL8* (L) were measured. *SLFN11*^-/-^ HEK293 cells [(D to F) and (J to L)], *SLFN11*^-/-^ DU145 cells (G), and *SLFN11*^-/-^ 293A cells [(H) and (I)] were generated by CRISPRLJCas9. pcDNA3.1-SLFN11 was transfected for complementary expression (F). The results of qPCR assays [(E to I) and (L)] are expressed as the mean ± s.d. from three technical replicates, and the data are representative of three independent experiments (D to K).

Additionally, the exhibited high immunostimulatory activity of AAV2 gDNA supports the recent hypothesis that the accumulation of AAV2 gDNA could lead to immune-related hepatic disease (*7*). Furthermore, adhering to this motif rule, we searched for and identified stimulatory ssDNA minimal mimics from viral genomic ssDNA (Fig. 3H and table S2). These results suggest that the immunostimulatory potency of ssDNA is determined by its stimulatory motifs, regardless of the species of its origin.

TLR9 recognizes only unmethylated CpG ODNs and exhibits tolerance to the phosphorothioate (PS) backbone (*4*). However, in the case of intracellular stimulatory ODNs, the PS backbone disrupts immunostimulatory activities, while methylation of CGT/A motifs has little effect on their potency (fig. S4, J and K). Thus, although intracellular stimulatory ODNs share some sequence similarity with canonical TLR9-sensed ODNs (fig. S4I), they exhibit markedly different tolerances for these types of modifications. As a result, canonical TLR9-sensed CpG ODNs were not as potent as ODN-H when delivered into cells. Meanwhile, TLR9 deficiency did not affect the stimulatory activity of ODN-H (fig. S4L). Taken together, these elements define immunostimulatory motifs for intracellular ssDNA: CGT/A motifs with phosphodiester (PO) linkages and methylation tolerance, followed by poly G in the 3′ downstream. For simplicity, we refer to these stimulatory motifs as CGT motifs. These results further suggest the existence of a novel receptor for intracellular ssDNA.

### SLFN11 is essential for intracellular ssDNA-induced immune responses

To identify the potential receptors for intracellular ssDNA, we performed an unbiased genome-wide CRISPRLJCas9 screen (Fig. 3A). We focused on hit genes whose knockout abrogated ODN60-induced cell depletion. Intriguingly, SLFN11, a member of SLFN protein family, was most significantly enriched in both single-round and three-round screens, while other identified DNA sensors (*1, 2*), such as TLR9, AIM2, and cGAS, did not show high levels of enrichment (Fig. 3, B and C). SLFN11 was previously found to play roles in inhibiting HIV translation (*27*) and the tumor response to DNA-damaging agents (*28, 29*). To validate the role of SLFN11 in intracellular ssDNA-induced immune responses, we generated SLFN11-knockout clones from HEK293, 293A, and DU145 cells. The absence of SLFN11 completely blocked all previously identified immune responses, including the cytokine activation, decline in cell viability, and pathway activation that were triggered by ODN60 (fig. S5, A to C) and other top-ranked ODNs (Fig. 3, D and E and fig. S5, D and E). Complementary expression of SLFN11 in *SLFN11^-/-^*HEK293 cells restored their responsiveness to ODN60 (Fig. 3F and fig. S5F), although this complementary expression itself also automatically provoked partial immune responses. The robust blocking effects observed from *SLFN11* knockout were exclusively for CGT ODNs and not for other types of stimuli (Fig. 3G). Additionally, only cells bearing SLFN11 expression were responsive to CGT ODNs, and Hela cells and 293T cells do not express SLFN11 (fig. S5G and Fig. 1, F and G). These results consistently indicate that SLFN11 is essential for CGT ODN-induced immune responses.

Moreover, CGT ODNs derived from viral genomic sequences and the single-stranded gDNA of *E.coli*, AAV2, and HIV1 activated *TNF* and *CXCL8* expression exclusively in WT 293A cells, but not in *SLFN11^-/-^* 293A cells (Fig. 3, H and I). Importantly, SLFN11 depletion also impeded the HU-induced activation of cytokines without affecting the accumulation of endogenous ssDNA (Fig. 3, J to L). These results indicate that SLFN11 is necessary for the stimulatory effects of both endogenous and pathogen-derived ssDNA.

### SLFN11 is a CGT ssDNA-binding protein

To determine whether SLFN11 could recognize the CGT motif and bind stimulatory ssDNA, we developed a biotin-pulldown complementary method named luminous ODN immunoprecipitation (ODIP) to measure the intracellular ODN-binding activity of SLFNs (Fig. 4A). ODN60, but not ODN60rc, was specifically immunoprecipitated with SLFN11, and the ODIP signal of ODN60 was only detected in cells expressing SLFN11 (fig. S6, A and B), suggesting that the ODIP assay is both specific and reliable. Notably, the binding of ODN to SLFN11 was dependent on CGT motif, as any single mutation in the CGT motif of ODN60 dramatically reduced its binding with SLFN11 (Fig. 4, B and C), while the gain of a CGT motif in ODN60rc enhanced its binding with SLFN11 (fig. S6C). In line with this, SLFN11 specifically bound to top-ranked ODNs but not bottom-ranked ODNs (Fig. 4, D and E). Collectively, these results suggest that the presence of CGT motifs in ssDNA enables a high affinity for SLFN11, which in turn determines the immunostimulatory properties of intracellular ssDNA bearing these motifs.

**Fig. 4.**
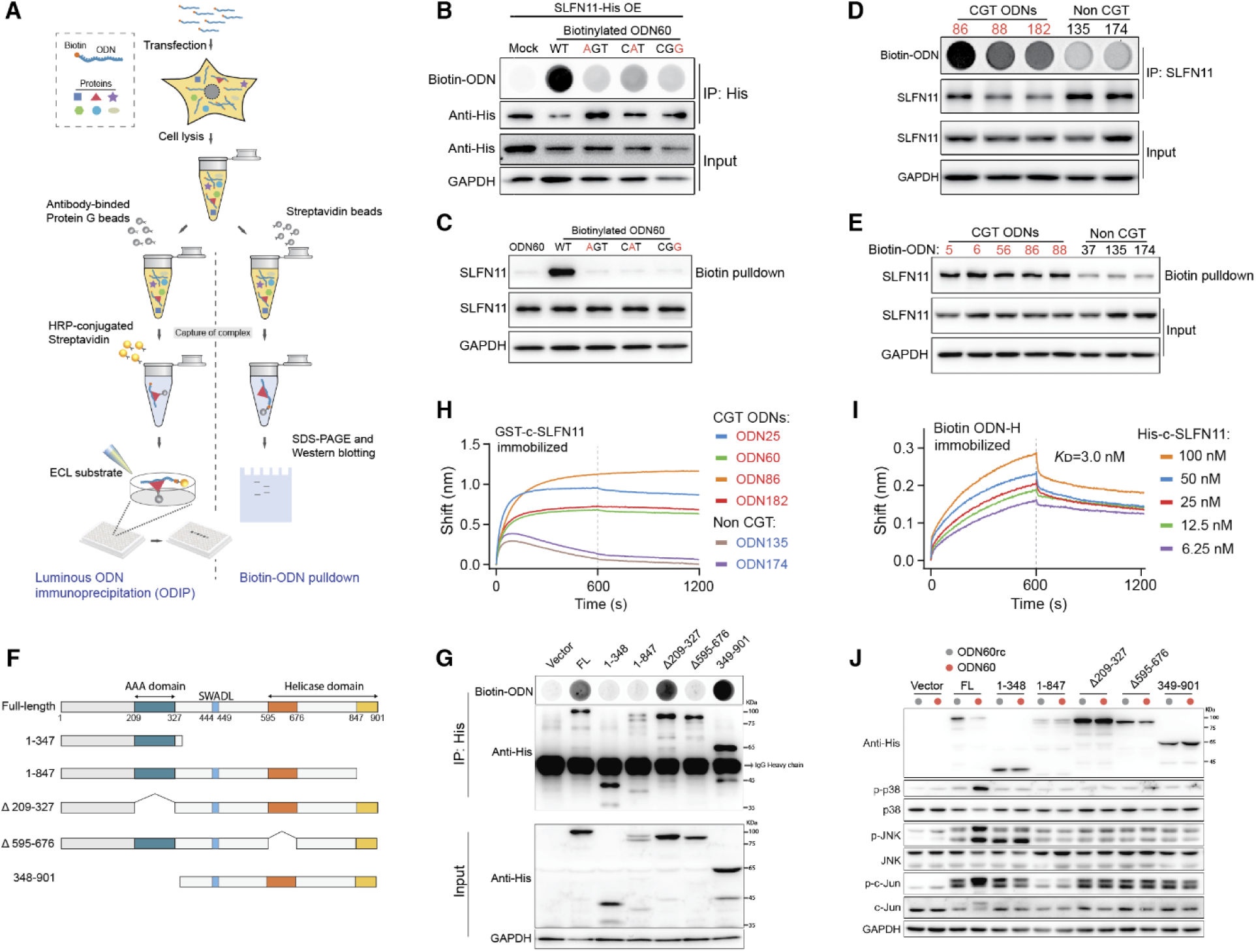
SLFN11 binds ssDNA containing a CGT motif. (**A**) Diagram of the luminous ODN immunoprecipitation (ODIP) and biotin-ODN pulldown assays. Cells were transfected with biotinylated ODN and lysed 6∼12 hours later. For ODIP, protein A/G-coated beads attached to a primary antibody were used to immunoprecipitate the protein and its bound ODNs from the cell lysate. The immunoprecipitated biotin-ODNs were then quantified using streptavidin (SA)- conjugated HRP to detect the enhanced chemiluminescence (ECL) substrate. For biotin-ODN pulldown assays, SA-coupled beads were used to pull down the biotin-ODN and the bound protein from the cell lysate. The captured proteins were analysed via immunoblotting. (**B** to **E**) The binding of SLFN11 to the indicated ODNs was analysed by ODIP [(B) and (D)] and biotin-ODN pulldown assays [(C) and (E)]. Biotinylated ODN60 or the indicated ODN60 mutants [(B) and (C)], or the indicated CGT ODNs and control ODNs [(D) and (E)] were transfected into HEK293 cells (C to E) or *SLFN11*^-/-^ HEK293 cells with re-expressed His-tag SLFN11 (B). Mock, transfection regent. (**F** to **J**) Full-length SLFN11 and the indicated truncated forms of SLFN11 (F) were re-expressed in *SLFN11*^-/-^ HEK293 cells, which were subsequently transfected with biotinylated ODN60. The binding activity of truncated SLFN11 with ODN60 was analysed by ODIP (G), and the ability of the truncated constructs of SLFN11 to rescue ODN-induced pathway activation was analysed by immunoblotting (J). (**H** and **I**) BioLayer Interferometry (BLI) assays of SLFN11 binding activity with CGT ODNs and control ODNs (H) and ODN-H (I). GST-tagged (H) and His-tagged (I) C-terminal SLFN11 (residues 349-901) constructs were expressed and purified from *E. coli* BL21 (DE3) cells and Sf9 cells, respectively. Sensorgrams reflecting the binding to ODN-H with gradient concentrations of His-tagged C-terminal SLFN11 and the calculated dissociation constants (K_D_) were calculated (I). All data shown are representative of three independent experiments.

SLFN11 belongs to subgroup III of the SLFN family, the members of which contain three predicted domains (*30, 31*): the N-terminal AAA domain, SWADL domain, and C-terminal helicase domain (fig. S7A). Our truncation analyses revealed that the C-terminal fragment of SLFN11 (residues 349 to 901) exhibited ssDNA binding capability, while the N-terminal fragment (residues 1 to 348) did not. Moreover, any indicated deletions within residues 349 to 901 significantly diminished the ssDNA-binding activity of SLFN11 (Fig. 4, F and G). The direct ssDNA-binding activity of SLFN11 was further confirmed using purified GST-tagged and His-tagged C-terminal SLFN11 (Fig 4, H and I and fig. S6, D to G). The calculated dissociation constants (*K*_d_) between SLFN11 and CGT ODNs were 3.0∼4.0 nM. During the preparation of this manuscript, another group also reported the ssDNA binding activity of SLFN11, but without exploring its biological significance (*32*). Notably, only full-length SLFN11, but not any truncated forms, was capable of restoring responsiveness to ODN60 in *SLFN11^-/-^* HEK293 cells (Fig. 4J). This suggested that while only the C-terminal helicase domain is responsible for ssDNA binding, the N-terminal domain is also required for signaling the presence of intracellular ssDNA. Collectively, the high binding affinity of SLFN11 for CGT ODNs, coupled with its essential role in intracellular ssDNA-induced immune responses, strongly suggests that SLFN11 functions as an intracellular ssDNA immune sensor.

Compared with other SLFN subgroups, only subgroup III that possesses the complete C-terminal ssDNA-binding domain is conserved from elephant fish to humans (fig. S7E). In addition, the subgroup III SLFNs display a similar evolutionary distribution among taxa to that of dsDNA sensors (fig. S7F). These data suggest that the C-terminal ssDNA binding domains of SLFNs are subject to extensive selective pressure. However, of all subgroup III SLFNs expressed in humans (fig. S7, C to D), only SLFN11 displayed detectable ssDNA binding activity (fig. S6H), consistent with our CRISPRLJCas9 screen data.

### Cytosolic translocation of ssDNA-bound SLFN11

SLFN11 has been reported to be a nuclear protein (*28, 29, 33*), as determined by immunofluorescence. Using the same detection method, we confirmed this finding by observing a diffuse nuclear distribution of SLFN11 under basal conditions. However, upon stimulation with CGT ODNs, SLFN11 was exported out of the nucleus and aggregated into puncta in the cytoplasm, colocalizing with CGT ODNs (fig. S8, A to C). This CGT ODN-induced translocation was further confirmed by live-cell imaging of *SLFN11^-/-^*cells with restored expression of SLFN11-GFP (movie S2). When examining cytoplasmic and nuclear extracts by immunoblotting, we found that under basal conditions, the distribution of endogenous SLFN11 was influenced by different extraction protocols (fig. S8D). This discrepancy may arise from the varying efficiency of isolating the perinuclear region in each protocol. To address this, we employed a cellular dissection technique that allowed for the compartmentalization of cells into three fractions: nucleus, perinucleus, and cytosol (*34*). SLFN11 was predominantly found in the perinuclear compartment under basal conditions, but translocated to the cytosolic compartment upon CGT ODN stimulation (fig. S8, E and F). This led us to hypothesize that the binding of CGT ODNs to SLFN11 may induce the cytosolic translocation. To test this hypothesis, we conducted biotin-ODN pulldown and native ODN PAGE assay. These results revealed that cytosolic translocated SLFN11 was indeed bound by CGT ODNs (fig. S8, G and H), consistent with the immunofluorescence observations that SLFN11 and CGT ODNs were colocalized in the cytoplasm (fig. S8, A to C). In addition to HEK293 cells, CGT ODNs also triggered cytosolic translocation of SLFN11 and colocalized with cytosolic SLFN11 in DU145 cells (fig. S8, I to K). These findings suggest that SLFN11 can translocate to the cytoplasm following physical recognition of ssDNA, further reinforcing its role as a sensor of intracellular ssDNA.

### SLFN9, the homologue of SLFN11, functions as the immune sensor for intracellular ssDNA in mice

The murine genome lacks *SLFN11* (fig. S7B). To further investigate the in vivo role of SLFNs as intracellular ssDNA sensors, we screened murine SLFNs that could functionally complement human SLFN11 in *SLFN11^-/-^* HEK293 cells. Among all murine subgroup III SLFNs, which are homologous to human SLFN11 (fig. S7, B and C), only SLFN9 exhibited detectable ssDNA binding ability and was able to restore ssDNA presence signaling (fig. S6, I and J). Therefore, we generated *Slfn9*-knockout mice (fig. S9A). Initial assessment of the *Slfn9^-/-^*mice revealed that they are healthy and fertile in our colony under specific-pathogen-free conditions.

We initially investigated the role of SLFN9 in response to stimulatory ssDNA in primary cells. In primary bone-marrow-derived macrophages (BMDMs), *Slfn9* deficiency blocked the activation of *Ifnb1*, *Il6,* and *Cxcl2* by stimulatory CGT ODNs, ODN-H and HIV6441 (Fig. 3H), but not by poly (dA:dT), poly (I:C) or LPS (fig. S9B), suggesting that SLFN9 was specifically required for CGT ODN-induced responses. Since HIV6441 is more potent than ODN-H in murine cells, we selected HIV6441 as a representive CGT ODNs for the subsequent experiments in mice. To explore the roles of SLFN9 and TLR9 in response to different types of ODNs, we transfected HIV6441 and all three types of canonical TLR9-sensed CpG ODNs (fig. S4I), ODN1585, ODN1826, and ODN2395, into primary mouse adult fibroblasts (MAFs), lung fibroblasts (LFs), primary BMDMs, and plasmacytoid dendritic cells (pDCs), respectively. While all types ODNs stimulated IFNα production in WT and *Slfn9^-/-^* pDCs but not in *Tlr9^-/-^* pDCs, in consistent with the reported role of TLR9 in pDCs (*4*), only the knockout of *Slfn9*, but not that of *Tlr9*, blocked HIV6441-induced *Ifnb1* expression in MAFs, LFs, and BMDMs (fig. S9, C and D). In contrast, none of the canonical CpG ODNs exhibited a stimulatory potency comparable to HIV6441 in these cell types. These results suggest that canonical CpG ODNs are much less effective than CGT ODNs in activating SLFN9 intracellularly. Similar to SLFN11, SLFN9 plays a crucial role in sensing intracellular ssDNA across various cell types, whereas TLR9 does not possess this broad functionality.

Next, we extended our findings from primary murine cells to in vivo level by administering WT and *Slfn9^-/-^* mice with CGT ssDNA delivered intracellularly using the lipid nanoparticle (LNP) system (*35*), which was also utilized in the SARS-CoV-2 mRNA vaccine. Since intravenous injection of the LNPs primarily resulted in intracellular delivery to the liver (*35*) (Fig. 5A), we focused on assessing the role of SLFN9 in the hepatic response to the ODNs. In neonatal WT mice, injection of CGT ODN-LNPs led to activation of cytokine expression and the STAT1, JNK, and NF-κB pathways in the liver (Fig. 5, B and C), while empty LNPs did not evoke these responses. Absence of *Slfn9* almost completely abolished all CGT ODN-induced responses. Similarly, in adult mice, *Slfn9* was also required for CGT ODN-LNPs to trigger the expression of cytokines (Fig. 5D), except for *Ifnb1*. It is commonly believed that the injection of CpG ODNs into mice is nontoxic, even at quite high doses (*36*). However, we found that intravenous administration of CGT ODN-LNPs (3 mg/kg) was sufficient to cause extensive hepatocyte necrosis and lobular architecture disruption in the liver, along with elevated serum levels of alanine aminotransferase (ALT) and aspartate aminotransferase (AST) (Fig. 5, E and F). Septic shock was observed in mice exposed to 5 mg/kg of CGT ODN-LNPs (Fig. 5G). Notably, these stimulatory effects were observed only in WT mice and not in *Slfn9^-/-^*mice. We have identified that AAV2 gDNA induces SLFN11-dependent immune responses in human cells (Fig. 3I), and AAV2 gDNA is linked to unexplained acute hepatitis in children (*7*). Therefore, we injected AAV2 gDNA encompassed by LNPs into WT and *Slfn9^-/-^* juvenile mice. Consistently, SLFN9 is also required for AAV2 gDNA-LNPs induced sever immune responses (Fig. 5, H and I). Altogether, these in vivo results suggest that intracellular accumulation of stimulatory ssDNA can induce inflammation and hepatic necrosis and cause hepatitis in a SLFN9-dependent manner. Given the resemblance of these phenotypes to the AAV-induced severe hepatotoxicity observed during gene therapy and infection (*6, 7, 37*), SLFN11 may play a critical role in mediating AAV-induced immune responses in human.

**Fig. 5.**
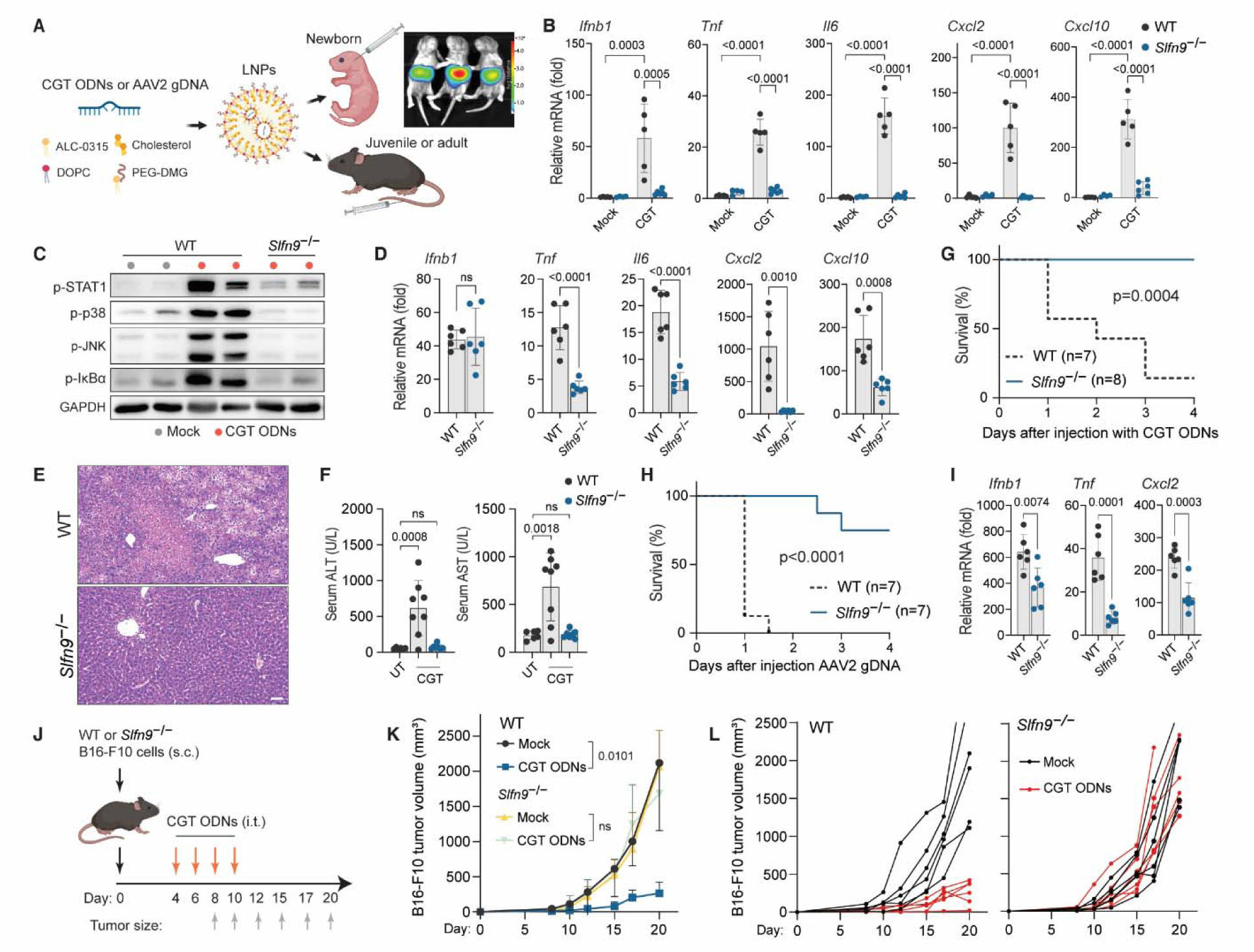
SLFN9 is essential for intracellular ssDNA-induced immune responses in mice. (**A**) Schematic illustration of the intravenous injection of ODNs or AAV2 gDNA encapsulated in LNPs; In vivo bioluminescence showing tissue distribution of intravenously injected luciferase mRNA encapsulated by LNPs. (**B** and **C**) WT and *Slfn9*^-/-^ newborn mice were injected with 3 mg/kg CGT ODN-LNPs (WT pups, *n* = 5 each treatment; *Slfn9*^-/-^ pups, *n* = 4 for mock and *n* = 6 for CGT ODN-LNPs), and liver samples were collected 6 hours post-injection. The mock groups were injected with empty LNPs. Cytokine levels (B) and immune pathway activation (C) were evaluated by qPCR and immunoblotting, respectively. (**D** to **F**) Adult WT and *Slfn9*^-/-^ mice were intravenously injected with 3 mg/kg CGT ODN-LNPs. The indicated cytokine expression levels were examined by qPCR (*n* = 6 each group) 6 hours post-injection (D); liver histological changes represented by Hematoxylin and Eosin (H&E) staining (E) and the ALT and AST levels in serum (F) were measured (WT mice, *n* = 6 for untreated control, *n* = 8 for CGT ODN-LNPs injection; *Slfn9*^-/-^ mice, *n* = 7 for CGT ODN-LNPs injection) at 3 days post-injection. Scale bar (E), 50 μm. (**G**) The survival of adult WT and *Slfn9*^-/-^ mice challenged with 5 mg/kg CGT ODN-LNPs was monitored over a span of 4 days. (**H** and **I**) Juvenile WT and *Slfn9*^-/-^ mice were intravenously injected with 3 mg/kg AAV2 gDNA-LNPs. The survival (H) and the indicated cytokine expression level (*n* = 6 each group) (I) were measured as in [(G) and (D)], respectively. (**J** to **L**) WT mice (*n* = 6 each group) were subcutaneously (s.c.) implanted with WT or *Slfn9*^-/-^ B16-F10 cells and then intratumorally (i.t.) injected with CGT ODNs encapsulated by transfection reagent. The treatment scheme is shown (J). The average tumor volumes of each group (K) and the tumor volumes of individual mice (L) at the indicated time points were measured. The data shown are representative of two (D), three [(E) to (L)] or four [(B) and (C)] independent experiments, with symbols indicating individual mice, and are shown as the mean ± s.d. Statistical significance was determined by unpaired two-tailed Student’s t test [(D) and (I)], one-way (F) and two-way [(B) and (K)] ANOVA with post hoc Bonferroni test and the log-rank test [(G) and (H)]; ns, not significant.

Previous basic and clinical studies have demonstrated the intratumoral monotherapy activity of CpG ODNs (*4, 38*), we therefore asked whether the superior potency of CGT ODNs in activating SLFN9 and inducing cell death could make them more suitable agents for cancer treatment. To explore this hypothesis, we generated *Slfn9^-/-^* and *Tlr9^-/-^* B16-F10 cells, respectively, even though WT B16-F10 cells express almost no *Tlr9* (fig. S9E). We found that the immunostimulatory activities of CGT ODNs observed in human cells were highly reproducible in B16-F10 cells, which were dependent on SLFN9 but not TLR9 (fig. S9, F to H). To further corroborate these findings in vivo, WT and *Slfn9^-/-^* B16-F10 cells were subcutaneously engrafted into mice (Fig. 5J). Consistent with previous results, intratumoral transfection of CGT ODNs significantly inhibited WT tumor growth but had no impact on the growth of *Slfn9^-/-^* tumors (Fig. 5, K and L), suggesting that SLFN9 is a crucial target in tumor cells for immunotherapy. Since Coley’s toxin are derived from heat-killed bacteria and bacterial ssDNA has been shown to be the major active component in Bacillus Calmette-Guerin (BCG) treatment for cancer (*4, 39, 40*), SLFN11/9 might also be activated in these strategies.

## Discussion

TLRs and intracellular surveillance systems have evolved to detect a similar range of PAMPs outside and inside cells, respectively. While TLR9, as a sensor for CpG ODNs, has been studied more than 20 years (*26*), the sensor responsible for detecting intracellular ssDNA remains unknown, largely due to the inconsistent stimulatory effects observed in previous studies involving the transfection of certain ODNs containing particular sequences (*3, 17, 19–22*). In this study, by utilizing genomic ssDNA and screening a short ssDNA library, we revealed that the immunostimulatory properties of intracellular ssDNA are highly sequence-dependent. Building on this insight, we identified the stimulatory motifs, developed the minimal stimulatory ssDNA mimics (CGT ODNs), characterized the immunostimulatory activities of intracellular ssDNA, and unequivocally identified SLFN11/9 as an intracellular surveillance system for ssDNA. These findings not only uncover a new dimension of the complementary relationship between TLRs and intracellular surveillance systems, but also broaden our understanding of the innate immune repertoire for detecting nucleic acids.

SLFN11 has previously been characterized as a nuclear protein with an N-terminal domain to cleavage tRNA (*28, 29, 32, 33, 41*). In this study, we have revealed a novel function of its C-terminal helicase domain in recognizing ssDNA containing a CGT motif, which leads to the translocation of SLFN11 to the cytosol. This suggests that ssDNA-activated SLFN11 may initiate downstream signaling events in the cytoplasm. However, additional studies will be required to fully elucidate the mechanism by which SLFN11 triggers cytokine expression and cell death. As the N-terminal of SLFN11 is also required for signaling the presence of ssDNA, it would be interesting to identify whether SLFN11/9 could function as CGT ssDNA-dependent RNases in the cytoplasm, which may suggest a new mode of immune activation. This study also predicts that the remaining C-terminal domains of subgroup III SLFNs may recognize other forms of nucleic acids or PAMPs.

As intracellular dsDNA surveillance systems have been well identified (*1, 42*), the majority of research on DNA-related diseases has been focused on dsDNA. Nevertheless, the intracellular accumulation of ssDNA is extensively observed during the pathogenesis of various diseases (*9–13*), such as tumor metastasis, senescence, and autoimmune diseases. Several proteins involved in dsDNA sensing, including cGAS (*10, 12*), IFI16 (*19*), and STING (*22*), have been reported to mediate intracellular ssDNA-induced type I interferons. Yet, our genome-wide CRISPRDCas9 screen did not significantly enrich these genes. Some cell types responsive to intracellular ssDNA even do not express these genes. Consistently, previous studies have demonstrated that cGAS and IFI16 are actually activated by double-stranded stretches of ssDNA, a dsDNA form, rather than the ssDNA structure itself (*18, 19, 42*), with STING functioning as a downstream factor of cGAS and IFI16 (*1, 17, 42*). In line with this, the binding affinity of cGAS and IFI16 to ssDNA was notably low, with respective *K*_d_ values of 1.5 µM (*1*) and 1.2 µM (*19*). This also implies that the immunostimulatory effects of ssDNA previously reported and attributed to these proteins might actually be stimulated by dsDNA (*10, 12, 18, 19, 22*). In contrast, SLFN11 binds stimulatory ssDNA with a much higher affinity (*K*_d_, 3∼4 nM). Hence, the identification of SLFN11/9 as the intracellular immune sensors establishes a currently unique mechanistical link between intracellular ssDNA and immune activation and diseases. Moreover, endogenous ssDNA-activated SLFN11 may serve as the mechanism connecting between DNA damage and innate immune responses (*43*). Interestingly, it has been reported that abundant ultrashort ssDNA derived from the mitochondria and the microbiota is present in serum (*25*). Considering a high proportion of SLFN11/9-sensed motifs in mtDNA and bacterial gDNA, potential aberrant activation of SLFN11 by these types of ssDNA could be relevant to numerous immune-related diseases.

The striking coincidence that both TLR9 and SLFN11/9 sense CpG-containing motifs implies the significance of CpG motifs for the immune system to discriminate between self and nonself DNA during evolution. However, since SLFN11/9 and TLR9 have distinct requirements regarding the surrounding sequences of CpG motifs and the structure of backbones, most commonly used CpG ODNs are not as effective as CGT ODNs in activating SLFN11/9. Additionally, as TLR9 is specifically expressed in B cells and pDCs, all the immune effects of ssDNA in human are thought to result from activating these cells (*4*). However, SLFN11/9 show a much broader expression profile across tissues, including tumor cells (*31, 44*). This considerably extends the range of ssDNA-responsive cells and tissues. Importantly, besides initiating cytokine expression, a primary activity of TLR9, SLFN11/9 can also trigger distinct immune responses, such as lytic cell death. Consequently, both TLR9 and SLFN11/9 sense ssDNA but may prompt different immune responses and outcomes. Taken together, the expanded range of responsive tissues and the novel immunostimulatory activities of intracellular ssDNA significantly reshape our mechanistic understanding of DNA-related diseases and therapeutic applications. For instance, while an association between the accumulation of AAV2 gDNA and a recent outbreak of acute hepatitis has been recognized (*7*), the causal link has remained elusive. Building on this new perspective, our study not only provides the first evidence that intracellular ssDNA can induce acute hepatitis, but also identifies SLFN11/9 as the mechanistic link. Therefore, our findings shed light on a previously overlooked facet of ssDNA immunity and underscore the significance of understanding CGT ssDNA as a novel type of stimulatory nucleic acids. Acting as intracellular ssDNA sensors, SLFN11/9 emerge as novel targets for developing CGT ODNs and small molecules to treat tumors and immune-related diseases and should be considered in ODN- or AAV-engaged therapeutic applications.

## Acknowledgments

We thank F. Shao (NIBS) for the long-term helpful discussions and expert comments on the manuscript, Z. Tian (USTC) for providing Tlr9 knockout mice, B. Zeng (ISMMS) and Q. Niu (Emory) for assistance to generate the ODN library, and Junko Murai (Keio) for the advice on SLFN11 immunofluorescence. We thank members of the Liu laboratory for helpful discussions and technical assistance.

## Author contributions

Conceptualization: P.Z.

Methodology: P.Z.

Investigation: P.Z., X.H., Z.L., Q.L., L.L., Y.J., S.L., X.Z., J.W.

Visualization: P.Z.

Funding acquisition: D.L.

Project administration: D.L., H.C., D.H., P.Z.

Supervision: D.L., H.C.

Writing – original draft: P.Z.

Writing – review & editing: P.Z., Z.L., D.L.

## Data and materials availability

All data are available in the main text or the supplementary materials. RNA-seq data used in this study have been deposited at the Sequence Read Archive database (PRJNA983563).

## Supplementary Materials

Materials and Methods

Figs. S1 to S9

Tables S1 to S2

References (*1–49*)

Movies S1 to S2

Data S1 to S2

## Supplemental Figure Legends

**Fig. S1.**
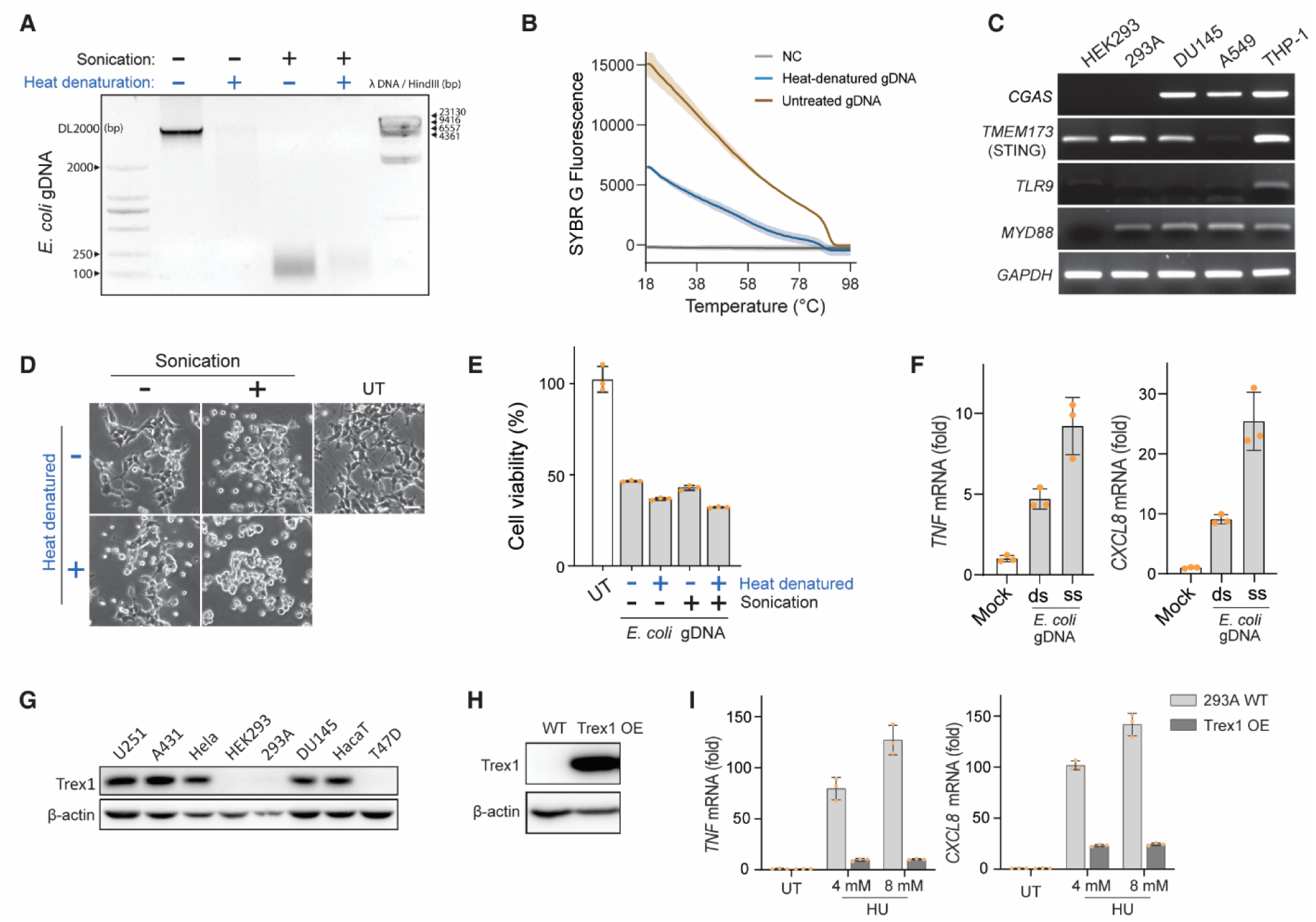
The *E. coli* genomic ssDNA and self ssDNA can activate immune responses intracellularly. (**A** and **B**) Sonicated and untreated *E. coli* gDNA was denatured by heat (98 °C for 5 min) to generate ssDNA. The resulting gDNA with the indicated treatments was resolved on a 1% agarose gel with ethidium bromide (EB) staining (A) and assessed by SYBR Green melting curve analysis (B), shaded regions indicate standard error; NC, SYBR Green without gDNA. (**C**) The expression levels of genes in cGAS or TLR9 signal pathway in the indicated cell types were analysed by RTLJPCR. The PCR products were resolved on a 1% agarose gel. (**D** to **F**) HEK293 cells were transfected with the indicated forms of *E. coli* gDNA. The ATP-based cell viability (D) and cell morphology (E) were examined 48 hours post-transfection, and *TNF* and *CXCL8* mRNA levels were measured by qPCR (F) 12 hours post-transfection. (**G** to **I**) Trex1 protein levels in the indicated cell lines (G) and Trex1-reconstituted HEK293 cells (H) were determined by immunoblotting. *TNF* and *CXCL8* mRNA levels (I) in WT and Trex-overexpressed (OE) HEK293 cells exposed to 4- or 8-mM HU were measured by qPCR. Data are shown as the mean ± s.d. from three technical replicates [(B), (E), (F), and (I)] and are representative of two [(A to C), (G), and (H)] or three [(D to F) and (I)] independent experiments.

**Fig. S2.**
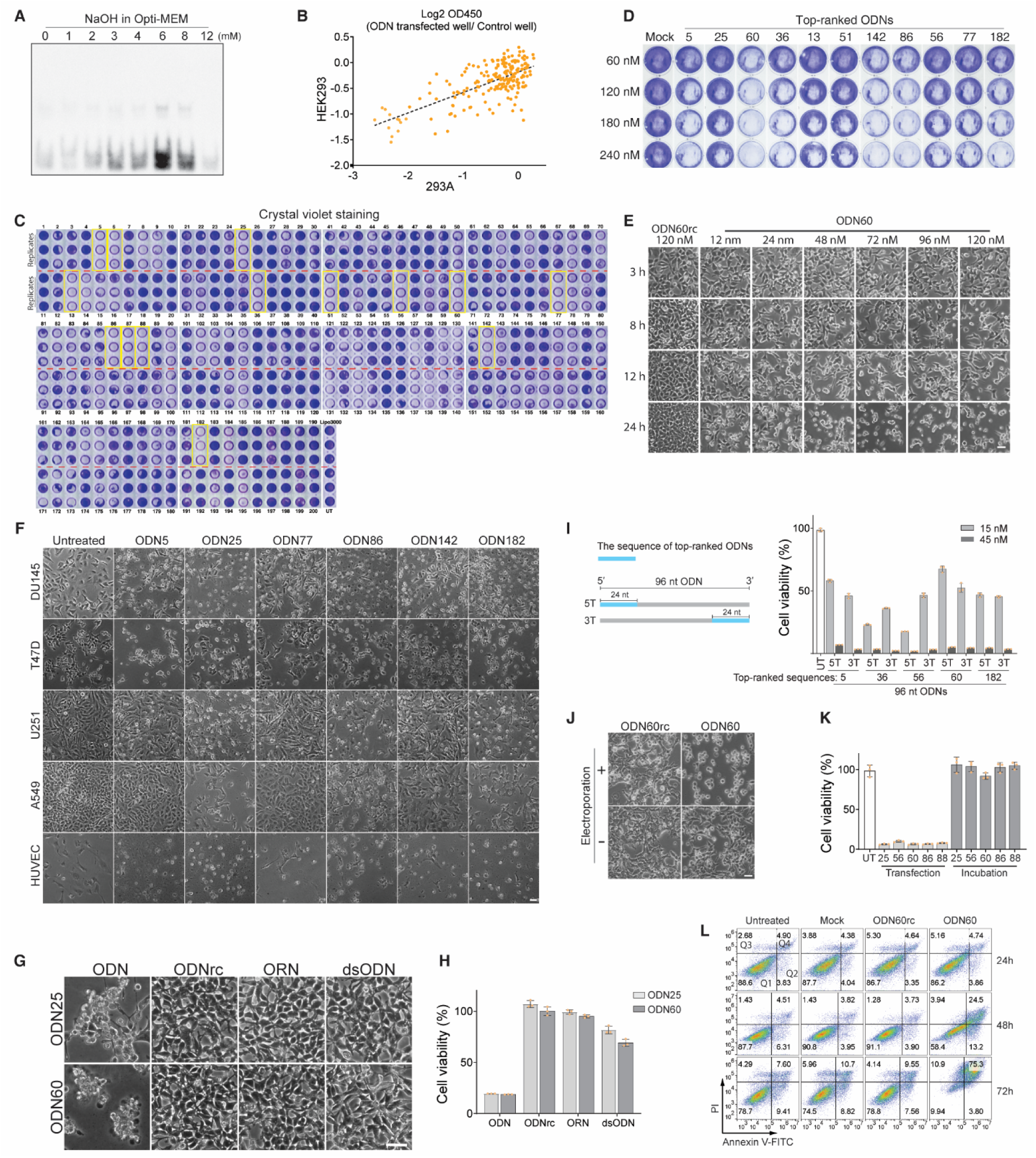
The decline in cell viability induced by intracellular ssDNA is a result of lytic cell death. (**A**) ODN transfection efficiency was sensitive to the pH of Opti-MEM. The biotinylated ODN harbouring the miR-148-3 sequence (tcagtgcactacagaactttgt) (500 ng) was transfected into HEK293 cells by Lipofectamine 3000 in Opti-MEM at the indicated NaOH concentrations. The ODN concentration in the lysates (6 hours post-transfection) were separated by 8% denaturing polyacrylamide gel and transferred onto a nylon membrane for detection using Streptavidin (SA)-HRP and a chemiluminescent substrate. (**B**) Correlation between CCK-8 assay-based cell viability of HEK293 and 293A cells for each ODN. (**C**) Crystal violet staining of 293A cells transfected with 200 ODNs in 96-well plates. Yellow rectangles indicate the top-ranked ODNs in Fig. 1B. (**D**) The dose response of the top-ranked ODNs was evaluated by crystal violet staining of attached HEK293 cells. (**E**) Time course and dose effect of ODN60 on cell morphology of transfected 293A cells. (**F**) Morphological changes in the indicated cell types 48 hours post-transfection with the top-ranked ODNs. (**G** and **H**) HEK293 cells transfected with the indicated forms of ODNs or ORNs were analysed for morphological changes (G) and ATP-based cell viability (H). ODNrc, reverse complementary ODNs; ORNs, oligoribonucleotides with the same sequence as the indicated ODN; dsODNs, double-stranded ODNs derived from annealing of ODN and ODNrc. (**I**) Effect of length on the ability of the top-ranked ODNs to decrease the cell viability of HEK293 cells. 5T/3T, 24-nt ODN sequence located in the 5′ or 3′ terminal of 96 nt ODNs. (**J**) Morphology of 293A cells 12 hours post-electroporation with ODN60 or ODN60rc. (**K**) HEK293 cells were transfected or extracellularly incubated with the indicated ODNs for 48 h, followed by ATP-based cell viability analysis. (**L**) Representative flow cytometry pseudocolor dot-plots of PI- and annexin V-stained HEK293 cells upon stimulation of transfection reagent (Mock), ODN60rc or ODN60. The ODNs were used at a concentration of 120 nM [(F to H) and (J to L)]. The scale bar [(D to G) and (J)], 50 μm. CCK-8 assay-based (B) and ATP-based [(H), (I), and (K)] cell viability is expressed as the mean or mean ± s.d. from three technical replicates. Data shown are representative of two [(B), (C), and (F)] or three [(A), (D), (E), and (G to L)] independent experiments.

**Fig. S3.**
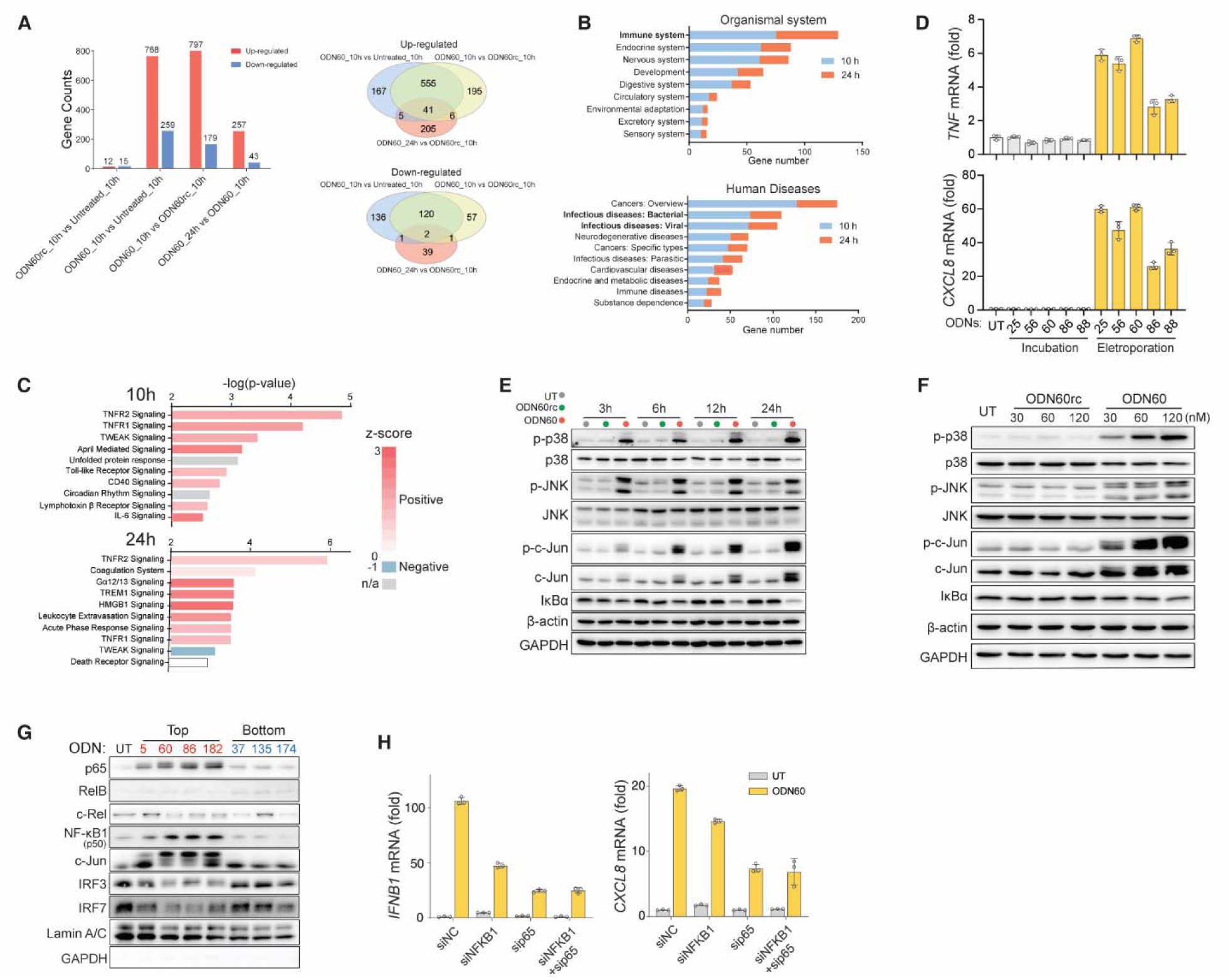
Intracellular ssDNA mimics are effective in activating cytokine expression and innate immune-related pathways. (**A t**o **C**) RNA-seq profiles of genes with upregulated or downregulated expression in 293A cells upon stimulation by ODN60 or ODN60rc (A). The differentially expressed genes (DEGs) represented in the Venn diagram (A) were enriched in KEGG pathways (B) and canonical pathways in the Ingenuity Pathway Analysis (IPA) database (C). (**D**) *TNF* and *CXCL8* expression levels were measured by qPCR in 293A cells electroporated or incubated with top-ranked ODNs for 6 hours. (**E** and **F**) The temporal (E) and dose (F) effects of ODN60 or ODN60rc on the activation of the indicated signal pathways in HEK293 cells. Cell lysates were examined by immunoblotting 12 hours post-transfection (F). (**G**) Nuclear fractions were isolated from 293A cells transfected with the indicated ODNs and analysed by immunoblotting with the indicated antibodies. (**H**) DU145 cells transfected with siRNA targeting negative control (siNC), NFKB1 (siNFKB1) or p65 (sip65) were stimulated with ODN60 for 12 hours. Subsequently, the expression levels of *TNF* and *CXCL8* were examined by qPCR. The ODNs were used at a concentration of 120 nM [(D), (E), (G), and(H)]. Data are shown as the mean ± s.d. [(D) and (H)] from three technical replicates and are representative of three independent experiments (D to H).

**Fig. S4.**
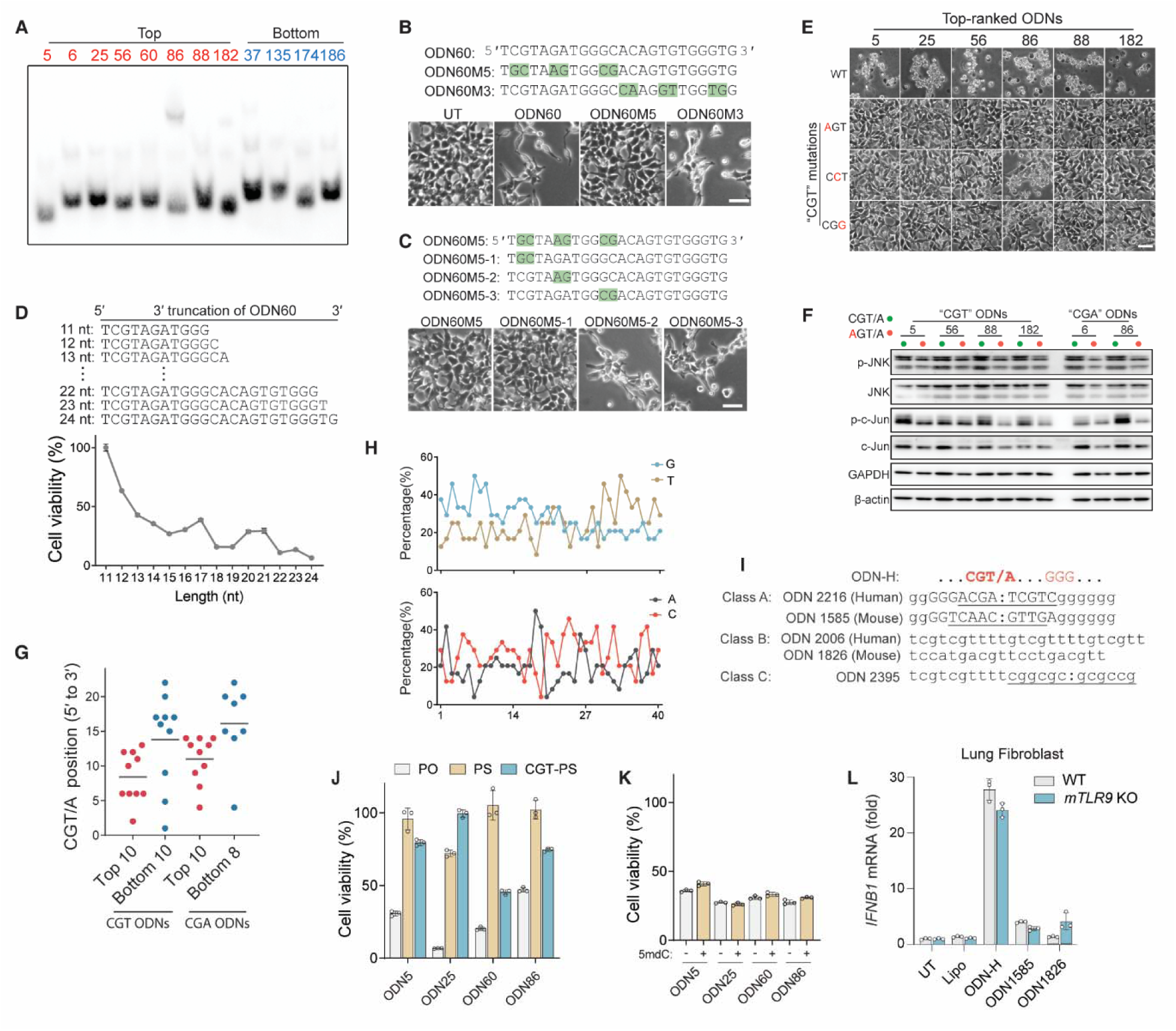
The immunostimulatory properties of ssDNA are dependent on a CGT motif. (**A**) Transfection efficiency of top- and bottom-ranked ODNs. HEK293 cells were transfected with the indicated biotinylated ODNs and analysed as described in fig. S2A. (**B** and **C**) Morphology changes of HEK293 cells transfected with ODN60 containing the exchange of the indicated adjacent nucleotides. (**D**) ATP-based cell viability of HEK293 cells transfected with ODN60 with 3′ terminal truncations. (**E** and **F**) Single-nucleotide sensitivity of CGT/A motif for the ability of the top-ranked ODNs to induce cell morphology changes (E) and activate JNK pathways (F). (**G**) Positions of the CGT/A motifs in top-ranked and bottom-ranked ODNs in Fig. 1E. (**H**) The percentage changes of four types of bases in CGT ODNs, relative to the stimulatory rank of Fig. 1E. (**I**) Sequence comparison of CGT ODNs with canonical CpG ODNs regarded as TLR9 agonists. Capital letters, phosphodiester (PO) link; lowercase case, phosphorothioate (PS) link; underline, palindromic sequence. (**J** and **K**) Effects of PS linkage (J) and 5-methyl-deoxy-cytidine modification (K) on the stimulatory potency of the top-ranked ODNs. (**L**) WT and TLR9-knockout mouse lung fibroblasts were transfected with the indicated ODNs, and *Ifnb1* mRNA expression was then measured by qPCR. HEK293 cells were transfected with the indicated ODNs at 120 mM, and phase-contrast images (scale bar, 50 μm) [(B), (C), and (E)] and ATP-based cell viability [(D), (J), and (K)] were examined 48 hours after transfection as representatives of immunostimulatory activity. Cell viability [(D), (J), and (K)] and qPCR (L) data are shown as the mean ± s.d. from three technical replicates. The data shown are representative of two (L) or three [(A to F), (J), and (K)] independent experiments.

**Fig. S5.**
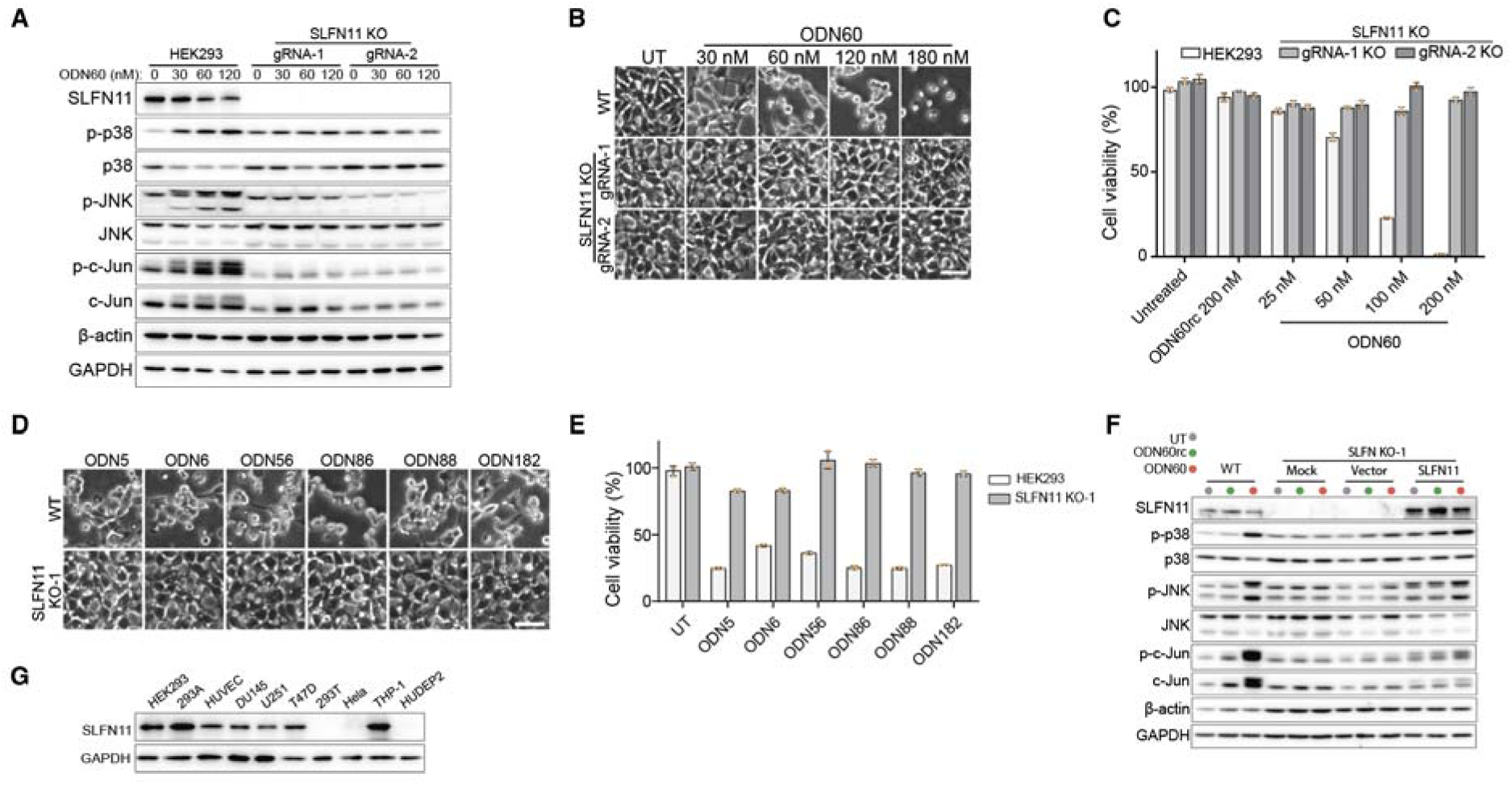
SLFN11 mediates intracellular ssDNA-induced immune responses. (**A** to **C**) Knockout of *SLFN11* affected ODN60-induced p38 and JNK activation (A), cell morphology changes (B) and cell viability decline (C). (**D** and **E**) cell morphology (D) and cell viability (E) were assessed in WT and *SLFN11^-/-^* HEK293 cells transfected with the top-ranked ODNs. (**F**) Effects of complementary expression of SLFN11 on ODN60-induced p38 and JNK activation, which was determined by immunoblotting. *SLFN11^-/-^* HEK293 cells were transfected with empty vector (pcDNA3.1) or pcDNA3.1-SLFN11. Mock, treated with only transfection reagent. (**G**) Cell lysates from the indicated types of cells were analysed by immunoblotting to determine the expression level of SLFN11. The ODNs were used at a concentration of 120 nM (D to F). Phase-contrast images of cells [(B) and (D)] with a scale bar (50 μm) and ATP-based cell viability [(C) and (E)] were analysed 48 hours post-transfection. ATP-based cell viability [(C) and (E)] results are expressed as the mean ± s.d. from three technical replicates. Data shown are representative of three independent experiments (A to G).

**Fig. S6.**
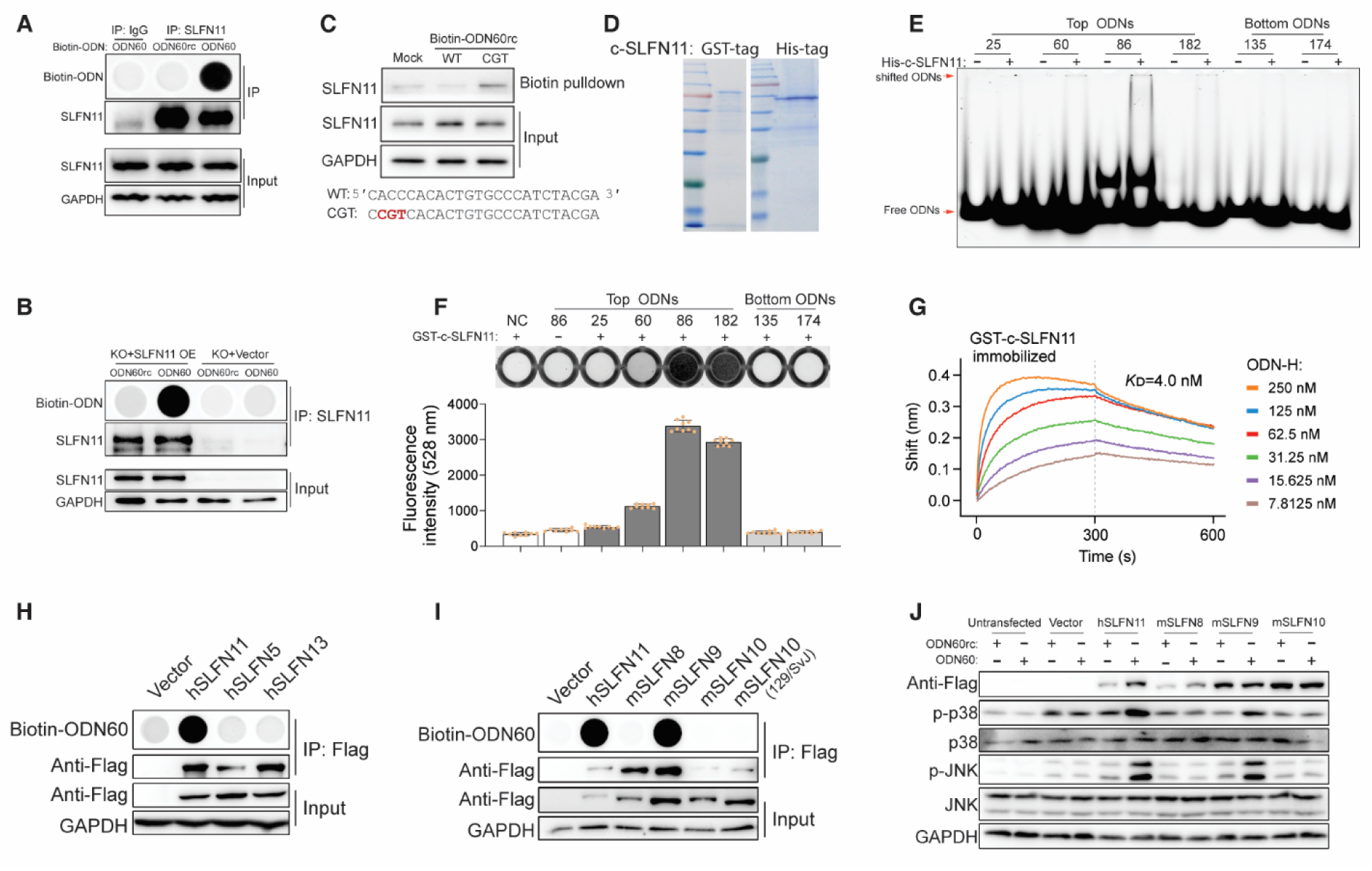
CGT ssDNA-binding activities of human and murine subgroup III SLFNs. (**A** and **B**) The binding activity of endogenous and re-expressed SLFN11 for ODN60 and ODN60rc was determined by ODIP after transfection of the indicated biotinylated ODNs into WT (A), SLFN11-deficient and SLFN11-restored (B) HEK293 cells. (**C**) The binding activity of ODN60rc or ODN60rc containing a CGT motif for SLFN11 were examined by biotin pull-down assay in HEK293 cells. (**D** to **G**) In vitro assays of the ODN-binding activity of purified SLFN11. GST-tagged and His-tagged SLFN11 C-terminal fragments (residues 349-901) were expressed and purified from *E. coli* BL21 (DE3) cells and Sf9 cells, respectively, and stained with Coomassie blue (D), Binding between the C-terminal fragment of SLFN11 and the indicated FAM-labeled ODNs was examined by electrophoretic mobility shift assay (EMSA) (E) and fluorescence ODN GST pulldown assay (F). The dissociation constant (K_D_) between GST-c- SLFN11 and ODN-H was measured using a BLI assay (G). (**H** to **J**) Restored expression of the indicated subgroup III SLFNs from humans (H) and mice (I) with an N terminal Flag tag in *SLFN11^-/-^*HEK293 cells. These cells were then transfected with biotinylated ODN60. The binding activity of SLFN proteins was determined by ODIP, and the ability of the indicated subgroup III SLFNs to restore the response to ODN60 was determined by immunoblotting (J). All data shown are representative of three independent experiments.

**Fig. S7.**
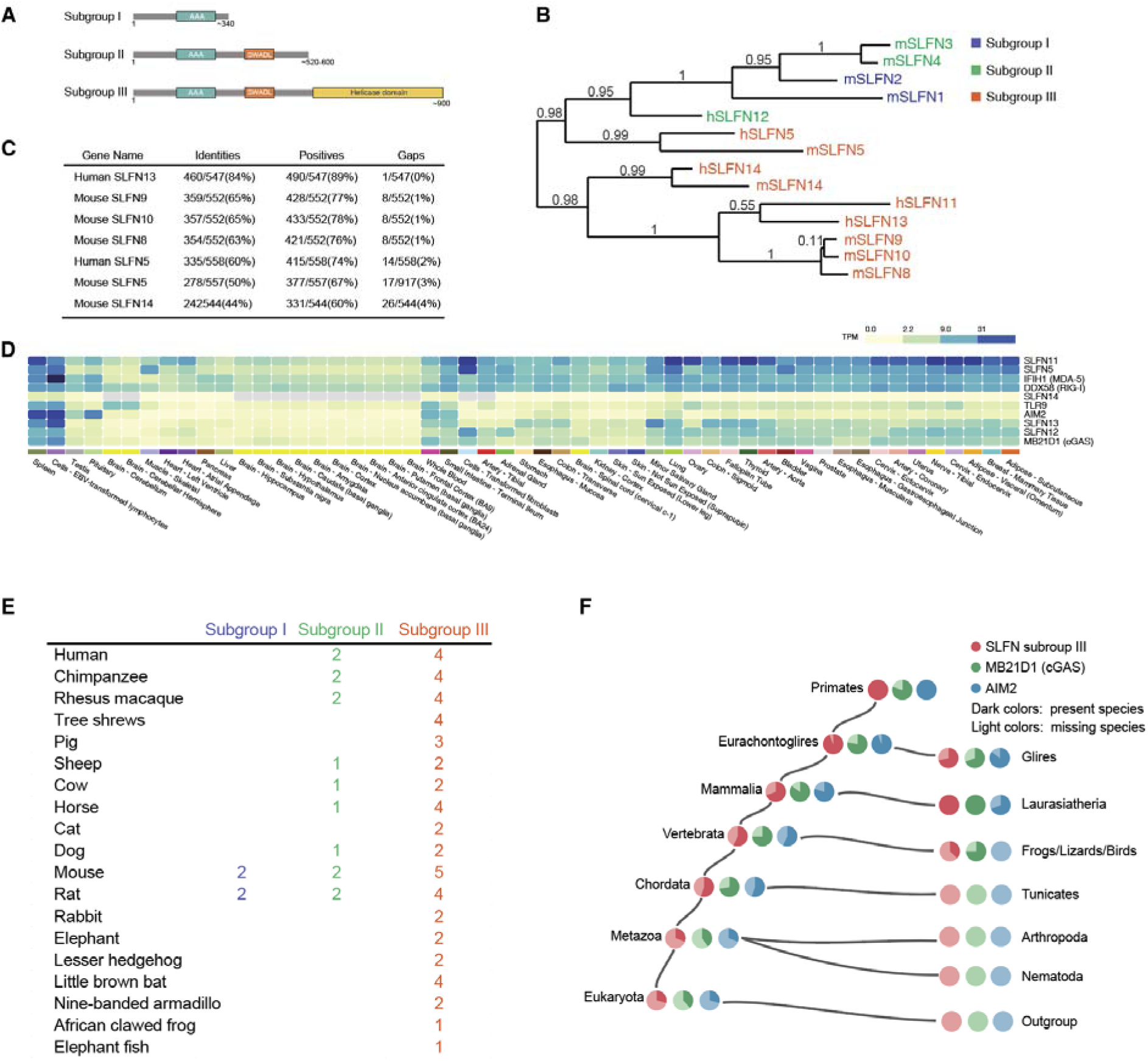
The SLFN family in humans and mice and its evolutionary tree. (**A** and **B**) The SLFN family is classified into three subgroups according to domain architecture (**A**). Phylogenetic tree of SLFNs in humans and mice, was generated using the Clustal Omega alignment and neighbor-joining method (**B**). (**C**) Proteins homologous to human SLFN11 (residues 349-901) were searched by protein-protein BLAST against the NCBI redundant protein sequence database for human and mouse proteins. (**D**) Expression levels of SLFN family proteins and identified RNA and dsDNA sensors in various human tissues from the Genotype-Tissue Expression (GTEx) database. (**E**) Distribution of SLFN subgroups among taxa based on the NCBI database. (**F**) Gene trees comparison of subgroup III SLFN, AIM2, and cGAS in the TreeFam 9 database.

**Fig. S8.**
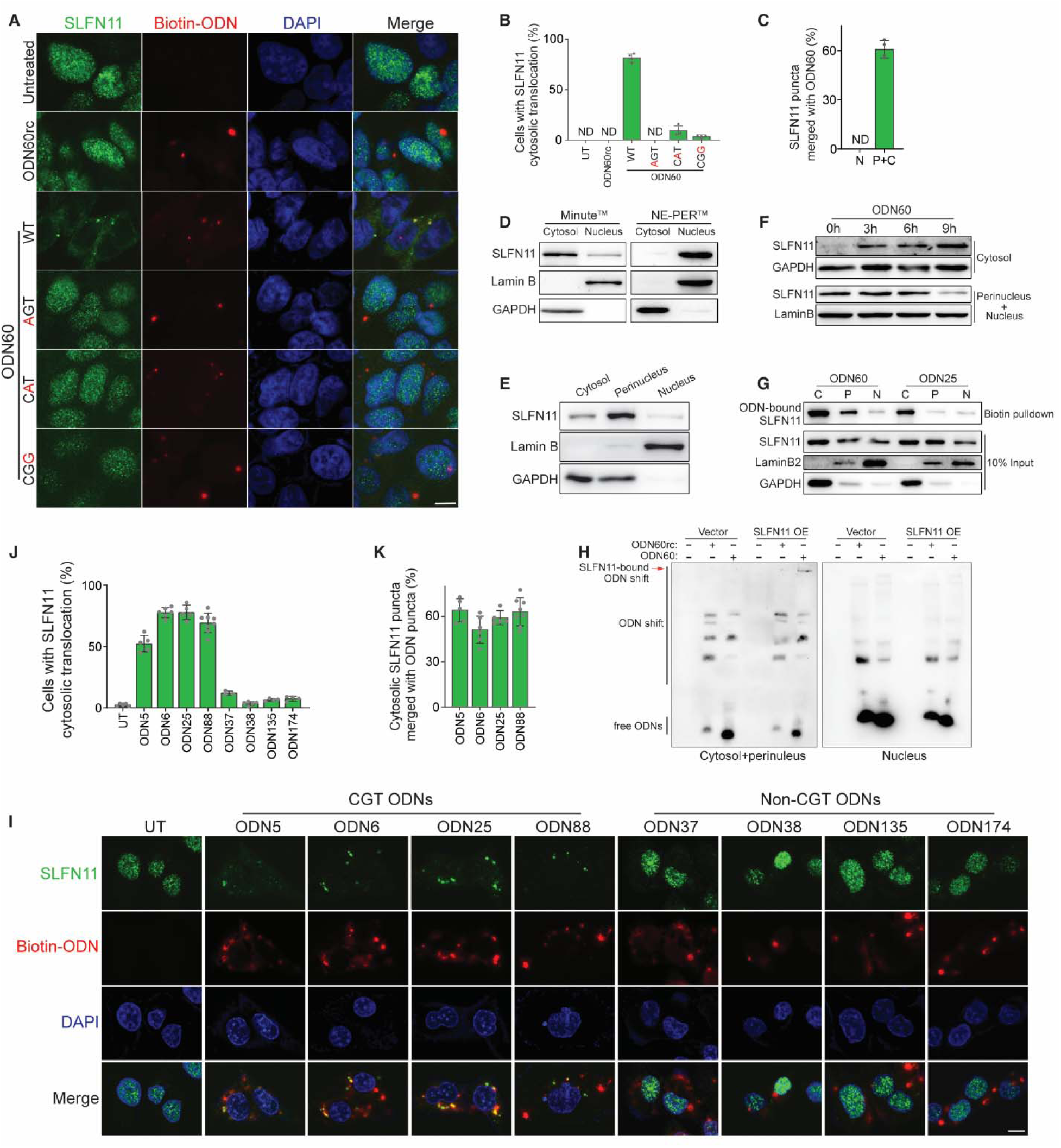
SLFN11 redistributes to the cytosol and colocalizes with CGT ODNs. (**A** to **C**) HEK293 cells were transfected with the indicated biotinylated ODNs and stained with SA-HRP to visualize biotinylated ODNs (red), antibody to visualize SLFN11 (green), and Hoechst 33342 to visualize nuclei (blue) 8 hours after transfection (A). The percentage of cells with SLFN11 cytosolic translocation (B) was quantified from 150 cells, and the percentage of SLFN11 puncta that had merged with ODN60 (C) was quantified from all SLFN11 puncta in these cells (n=4, mean ± s.d.). ND, not detectable; N, nucleus; P+C, perinucleus and cytosol. Scale bar (A), 10 μm. (**D** and **E**) HEK293 cells were fractionated into two (cytosol and nuclear) fractions (D) or three (cytosol, perinuclear, and nuclear) fractions (E). The SLFN11 distribution in the indicated fractions was determined by immunoblotting. Each fraction sample was adjusted to the same volume before loading. (**F**) The subcellular distribution of SLFN11 changed upon ODN60 stimulation. Nuclear-perinuclear and cytoplasmic fractions from HEK293 cells at the indicated time points after transfection with ODN60 were analysed by immunoblotting. (**G**) The subcellular location of ODN-bound SLFN11. Nuclear, perinuclear and cytosolic fractions were prepared from HEK293 cells transfected with biotin-ODN60 or biotin-ODN25, and the ODN-bound SLFN11 was then pulled down with SA beads and analysed by immunoblotting. The loading amounts of input and pulldown samples from three fractions were adjusted based on the SLFN11 levels in each input samples. N, nucleus; P, perinucleus; C, cytosol. (**H**) *SLFN11^-/-^* HEK293 cells with or without restored SLFN11 expression were transfected with biotinylated ODN60 or ODN60rc. The bound ODNs in the cytosolic-perinuclear and nuclear fractions were determined with native ODN PAGE. Each fraction sample was adjusted to the same volume before loading. (**I** to **K**) DU145 cells were transfected with the indicated biotinylated ODNs, fixed at 18 hours post-transfection, and then stained as in (B). Scale bar, 20 μm (I). The percentage of cells with cytosolic SLFN11 translocation (J) was quantified from 100 cells, and percentage of cytosolic SLFN11 puncta merged with indicated ODNs (K) was quantified from all cytosolic SLFN11 puncta in these cells (n=4∼8, mean ± s.d.). Data shown are representative of three [(A to C), (D to F), and (I to K)] or four [(G) and (H)] independent experiments.

**Fig. S9.**
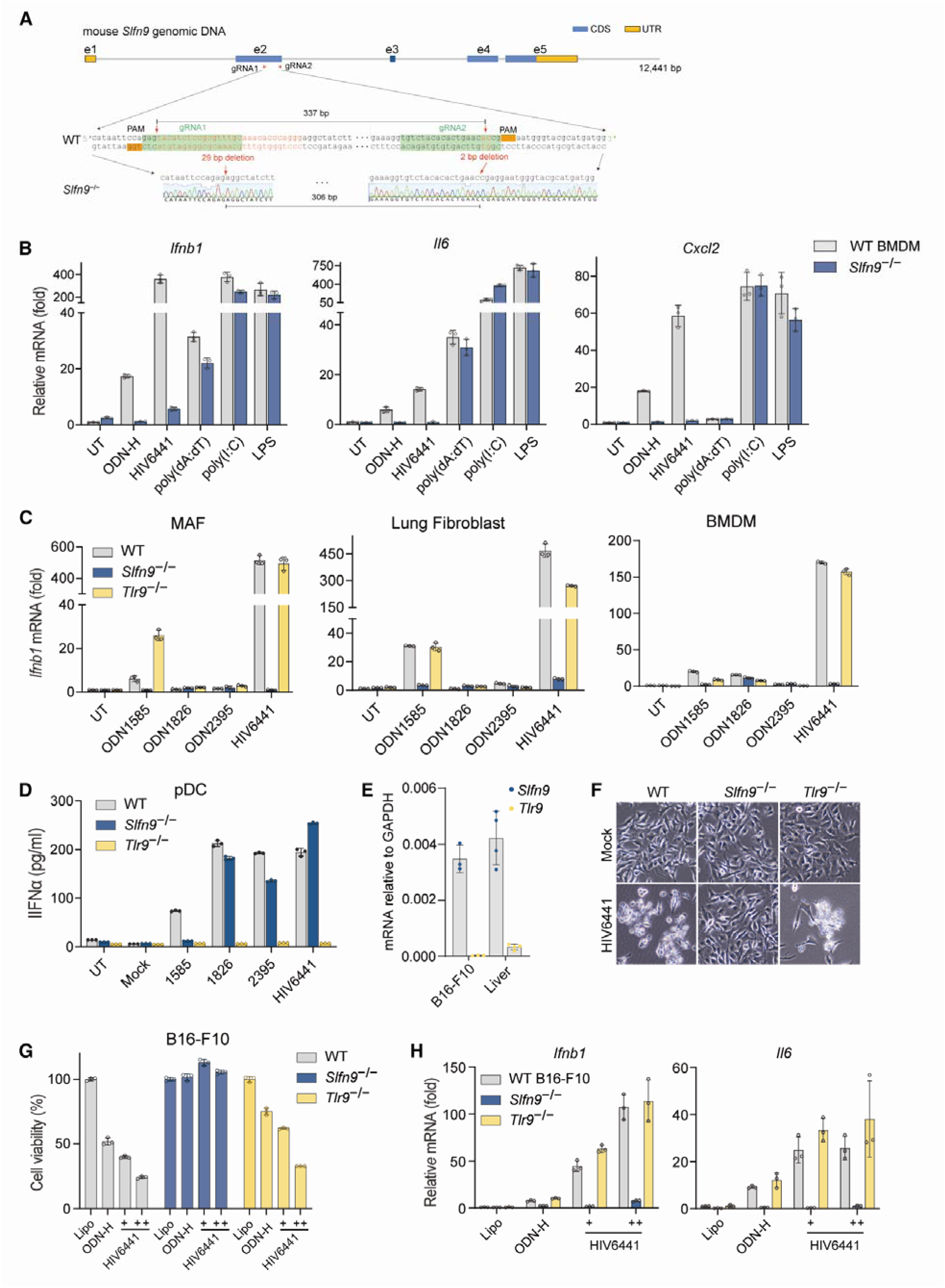
SLFN9 mediates CGT ODN-induced immune responses in mouse primary and cancer cells. (**A**) Characterization of *SLFN9*-KO mice generated by the microinjection of Cas9 mRNA and a pair of gRNAs targeting the *Slfn9* 2^nd^ exon into zygotes. The deleted nucleotides are highlighted in red and were identified by sequencing. (**B**) The expression levels of *Infb1*, *Il6* and *Cxcl2* in WT and *Slfn9*^-/-^ BMDMs 18 hours post-transfection with the indicated types of nucleotides or 4 hours post-stimulation with LPS were determined with qPCR. (**C** and **D**) The indicated primary cells from WT, *SLFN9*^-/-^ and *Trl9*^-/-^ mice were transfected with HIV6441 and the indicated canonical CpG ODNs. *Ifnb1* mRNA expression was determined by qPCR (C) and secreted IFNα levels were determined by ELISA (D). (**E**) *SLFN9* and *Trl9* expression level in B16-F10 and WT mouse livers were analysed by qPCR. (**F** to **H**) *SLFN9*^-/-^ and *Trl9*^-/-^ B16-F10 cells were generated and transfected with ODN-H and HIV6441, respectively. Changes in cell morphology (F), ATP-based cell viability (G) and the indicated cytokine expression (H) were examined. ODN-H and HIV6441 +, 1 ug/ml; HIV6441 ++, 2 ug/ml. Data are shown as the mean ± s.d. [(B to E), (G), and (H)] and representative of two [(C), (D), and (E)] and three [(B) and (F to H)] independent experiments.

## Materials and Methods

### Plasmids, antibodies and reagents

Complementary DNAs (cDNA) encoding human *SLFN11*, *SLFN13*, and *SLFN5* were amplified from reverse-transcribed cDNA of HEK293 cells; cDNAs encoding mouse *SLFN9* and *SLFN8* were amplified from reverse-transcribed cDNA of fetal liver of 14.5-day mouse embryos; cDNAs for mouse *SLFN10* were synthesized by Sangon Biotech (Shanghai). The desired gene cDNAs were inserted into pcDNA 3.1 vector with a C-terminal c-myc-6×His tag, pFlag-CMV vector with an N-terminal Flag tag or pEGFP vector with a C-terminal GFP tag. cDNA encoding *Trex1* was amplified from cDNA of DU145 cells and subcloned into pLV-EF1a-EGFP. All the truncated *SLFN11* coding sequences were amplified from full-length *SLFN11* cDNA and inserted into pcDNA 3.1 vector with a C-terminal c-myc tag and 6×His tag or pEGFP-C vector with a C-terminal fused GFP. For recombinant expression in *E. coli* and Sf9 cells, the coding region of *SLFN11* (349-901 residues) was cloned into pGEX-4T-1 and pFastbac1, respectively.

Antibodies against SLFN11 (D-2) (sc-515071), SLFN11 (E-4) (sc-374339), Trex1 (sc-271870) and LaminB2 (sc-377379) were obtained from Santa Cruz Biotechnology. Antibodies against phospho-p38 (#9211), p38 (#9212), phospho-JNK (#9251), JNK (#9252), c-Jun (#9165), phospho-c-Jun (Ser73) (#3270), phospho-NF-κB p65 (Ser536) (#3033), NF-κB p65 (#6956), RelB (#4922), cRel (#12707), NF-κB1 (#12540), NF-κB2 (#4882), IkBα (#9242), phospho-IkBα (#2859), IRF-3 (#4302), IRF-7 (#72073), Lamin A/C (#4777), phospho-STAT1 (#9171S), and rabbit anti-His (#12698) were purchased from Cell Signaling Technology. The other antibodies used in this study included rabbit anti-SLFN11 (Sigma, HPA023030), anti-ssDNA (Millipore, MAB3868), mouse anti-His (MBL, D291-3), anti-GFP (MBL, 598), anti-Flag (CMC-TAG, AT0022), anti-β-actin (Abcam, ab8227) and anti-GAPDH (ABclonal, AC002).

All ODNs and ORNs used in this study were synthesized by Invitrogen or Sango. Canonical CpG ODNs including ODN1585, ODN1826 and ODN2395 were from a mouse TLR9 agonist kit (Invivogen, tlrl-kit9m). Poly(I:C) (tlrl-pic), poly(dA:dT) (tlrl-patn), ISD (tlrl-isdn) and LPS (tlrl-peklps) were purchased from Invivogen. The bacterial genomic DNA used in this study was from *E. coli* strain B (Sigma, D4889). To knockdown NFKB1 and p65, the siRNAs were synthesized by JTS (Wuhan). All types of lipids used for nanoparticle formation were purchased from AVT (Shanghai) Pharmaceutical Tech Co., Ltd.

### Cell culture

HEK293, DU145, T47D, U251, A549 and B16-F10 cells and HUVEC were obtained from the Cell Resource Center of Peking Union Medical College (PUMC), which is the headquarter of National Science & Technology Infrastructure-National BioMedical Cell-Line Resource (NSTI-BMCR). Their species origins were confirmed with PCR. The identity of each cell line was authenticated with STR profiling (FBI, CODIS). All the results can be viewed on the website (http://cellresource.cn). The 293A cell line was purchased from Invitrogen. All cell lines were tested to be mycoplasma-negative by PCR test (YEASEN, 40601ES10). HEK293 and U251 cells were grown in Minimal Essential Medium with Earle’s Balanced Salts (MEM/EBSS), 293A cells were grown in Dulbecco’s Modified Eagle’s Medium (DMEM), DU145, T47D and B16-F10 cells were grown in RPMI 1640 medium (w/o Hepes), and A549 cells were grown in McCoy’s 5A (Modified) Medium. For primary cell culture, MAFs and LFs were cultured in DMEM; BMDMs and pDCs were cultured in RPMI-1640 medium containing 2 mM glutamine. All media were supplemented with 10% (vol/vol) fetal bovine serum and 1% Penicillin/Streptomycin/Amphotericin B (Solarbio, P7630). All cells were grown at 37°C in a 5% CO_2_ incubator. All types of medium were obtained from the Cell Resource Center of PUMC.

### Isolation of MAFs, LFs, BMDMs and pDCs

Primary MAFs were prepared from 6-week-old mice by following a standard procedure as previously described (*45*). Briefly, the mouse ear was cut and consequently digested with collagenase IV (Gibco, #17104019) (1,000 U/ml) and 0.05% trypsin. LFs and BMDMs were isolated from 6-week-old mice by following previously described procedures (*46*). To isolate primary LFs, lung tissues were cut and digested with 0.1% collagenase I and 0.2% trypsin. To obtain BMDMs, bone marrow cells collected from the femurs and tibiae of mice were cultured in RPMI-1640 medium containing 10% FBS, L-glutamine (2 mM) and mouse M-CSF (10 ng/ml) for 6 to 7 days. Mouse pDCs were isolated from 6-to-8-week-old mice using EasySep pDC isolation kit (STEMCELL, #18000). The spleens of the mice were disrupted in PBS and filtered through a 70 μm mesh nylon strainer. A total 1 × 10^8^ nucleated cells were used for pDC isolation by following the manufacturer’s protocol. The isolated pDCs were cultured in a 96-well plate and transfected with the indicated ODNs. The IFN-alpha secreted by pDCs was measured by an ELISA kit (R&D SYSTEM, 42120-1). All types of murine primary cells are on C57BL/6 genetic background.

### Single-stranded genomic DNA preparation

To shorten *E. coli* gDNA for transfection, the gDNA was sheared into ∼ 200-bp fragments using an S220 Focused-ultrasonicator (Covaris) according to the manufacturer’s protocol. The AAV2 genomic DNA (gDNA) was amplified from the pAAV2-RC plasmid, while the HIV gDNA was amplified from the psPAX2 plasmid using PCR. To prepare single-stranded gDNA, double-stranded gDNA was heat denatured in a boiling water bath for 10 min, and then immediately placed on ice.

### ODN library and viral CGT ODN generation

A genome-wide ODN library was generated according to the ClinVar VCF file (20170615 version) for GRCh37. Each missense site served as a seed site to randomly generate a 24-nt sequence according to the genome sequence. A total of 112,764 sequences were outputted, and 200 sequences among them were randomly picked for final ODN library synthesis and screening. For viral CGT ODN generation (Table S2), HIV2 (NC_001722), B19V (NC_000883) and PCV2 (MH931449) genome sequences were used to search the stimulatory motifs, CGT/A followed by a poly G. The codes will be publicly released at https://github.com/pengzhang-ssDNA/ssDNA-sensor.

### Analysis of proportion of stimulatory motifs in variety of gDNA

The gDNA sequences used for analysis of stimulatory motif proportion are available at the National Center for Biotechnology Information. The accession numbers: *E.coli* gDNA (NC_000913.3), HSV1 (NC_001806), Adv7 (AC_000018.1), HBV (X98077.1), HIV1 (NC_001802.1), HIV2 (NC_001722), B19V (NC_000883), AAV2 (AF043303), PCV1 (JN133302), PCV2 (MH931449), gorilla anellovirus (NC_030650), and human mtDNA (NC_012920.1). The sequences of spCas9 and saCas9 are from px330 and px601 plasmids, respectively. The codes will be publicly released at https://github.com/pengzhang-ssDNA/ssDNA-sensor.

### Nuclear acid transfection and electroporation

The day before transfection, cells were plated in 1 ml of growth medium so that they were 40-60% confluent at the time of transfection. gDNA or ODNs were transfected into cells with Lipofectamine 3000 (ThermoFisher, L3000001) according to the manufacturer’s protocol with some modifications. In detail, dilute gDNA or ODNs were diluted in Opti-MEM without P3000. The Lipofectamine-to-ODN/gDNA ratio was 1.5:1 (μl:μg). ODN transfection efficiency is sensitive to pH of Opti-MEM. As freshly-opened Opti-MEM is not suitable for ODNs transfection, we adjust the pH by adding 5 mM NaOH to a final concentration of 5-6 μM in Opti-MEM. To overexpress the indicated proteins, we used Vigofect (Vigorous, T001) for plasmid transfection into HEK293 and 293A cells based on the manufacturer’s instruction without modification. We transfected 1 μg of gDNA or plasmid or ODNs per 1 ml of culture medium (to a 120 nM final concentration in culture medium for 24 nt ODNs), unless otherwise noted. We electroporated 3 μg of ODNs or gDNA into 2 × 10^6^ HEK293 and 293A cells by Neon Transfection System (Thermo Fisher) based on the manufacturer’s introductions.

To perform siRNA knockdown in DU145 cells, 2 μl of 20 μM siRNA was incubated with 4 μl of INTERFERin (Polyplus Transfection) in 100 μl of Opti-MEM and then transfected into DU145 cells in a 12-well plate format. The culture medium was exchanged with fresh medium after 8 hours of transfection. Knockdown was performed for 48 hours before subsequent ODN transfection. siRNA sequences: siRELA (ggacauaugagaccuucaatt) and siNFKB1 (gcaaucauccaccuucauutt).

### Microscopy imaging of cell morphology

To examine cell morphology, cells were treated as indicated in the 12-well or 24-well plates (Corning) for static image capture or in glass-bottom 24-well plates (NEST) for live imaging. Phase contrast bright field microscopy were captured using Nikon TS100 or TS2 and processed in ImageJ. Live images were recorded with Lionheart FX or Cytation 5 (BioTek) and processed in Gen5 software. All imaging data shown are representative of at least three randomly selected fields.

### RNA-seq analysis and quantitative PCR

RNA was extracted from the indicated cells with TRIzol reagent (Invitrogen) and treated with DNase I. mRNA was enriched by Oligo(dT), fragmented and reverse-transcribed into cDNA libraries. The cDNA was then purified, end-repaired, adenylated and barcoded with multiplex adapters. PCR-amplified libraries were purified with AmpureXP beads and validated on the Agilent 2100 Bioanalyzer. After quantification by Qubit (Invitrogen), the normalized samples were sequenced using Illumina HiSeq X Ten. Before mapping, the reads were filtered to remove low-quality reads, adaptor-polluted reads and reads with a high content of unknown bases (N). The clean reads were then mapped to the human genome (GRCh37/hg19) using HISAT. Gene expression levels were calculated with RSEM. Genes with a log2 (fold change) > 2, P < 0.01 and FPKM >0.1 were deemed to be significantly differentially expressed between the two samples and were used for KEGG (www.kegg.jp) analysis and pathway functional enrichment analysis with QIAGEN’s Ingenuity tool (www.ingenuity.com).

For quantitative PCR (qPCR), extracted RNA from the cells with the indicated treatment was reverse-transcribed to cDNA with RevertAid First Strand cDNA Synthesis Kit (K1622, Thermo Scientific) according to the manufacturer’s protocol, with random hexamers. Quantitative PCR was performed with AceQ qPCR SYBR Green Master Mix (Q121, Vazyme) in an Eppendorf Realplex cycler.

### Cell viability and cell death assay

Cell viability was determined by the CellTiter-Glo Luminescent Cell Viability Assay (Promega, G7571), the Cell Counting Kit-8 (CCK-8) assay (Beyotime, C0042), or crystal violet staining (Beyotime, C0121). For the ATP-based cell viability assay, both adherent and suspended cells were collected and washed once with PBS. The cells were then resuspended in PBS for the cell viability assay. For the CCK-8 based cell viability assay, the CCK-8 reagent was directly added to the cell culture medium (ratio, 1:10 v/v) at the indicated time points. After a 30-min incubation period, the absorbance at 450 nm was measured using a microplate reader (Cytation5, BioTek). For cell death assay, the cells were stained by annexin v-FITC and PI (Beyotime, C1062) and measured by BD Accuri C6. All assays were performed according to the manufacturer’s instructions.

### Genome-wide CRISPRLJCas9 screening

A ready-to-use pooled lentiviral library (Brunello) for CRISPR screening was purchased from Beijing Genomtech. In the pilot experiment, the volume of the lentivirus library required to achieving an MOI of 0.3 for infecting 293A cell line was determined by puromycin selection in the 12-well plate format. For the large-scale screen, 3 × 10^7^ cells were seeded in four 15-cm dishes (20 ml media per dish) and 24 hours later, a total of 5 × 10^7^ cells were infected with lentivirus library. Twelve hours after infection, the cells were reseeded at a density of 1 × 10^7^ cells per 15-cm dish. Another 12 hours later, puromycin was added to a final concentration of 2 μg/ml and incubated for 36 to 48 hours. Approximately 1 × 10^8^ infected cells were divided into 3 groups: the untreated control, ODN60rc transfection and ODN60 transfection groups. For a single round of transfection, surviving cells were collected at 48 hours posttransfection. For three-round transfection, cells were stimulated by ODNs for 36 hours, seeded onto a new dish and recovery-cultured in fresh medium for 36 hours between rounds of transfection. The collected surviving cells were lysed in SNET buffer (20 nM Tris-HCL (pH 8.0), 5 mM EDTA, 400 mM NaCl, 1% SDS and 400 μg/ml Proteinase K) at 55 °C overnight. Proteinase K was inactivated by boiling the cell lysate for 10 min, and RNA was degraded with RNase. Chromosomal DNA was extracted by phenolDchloroform-isoamyl alcohol and precipitated by isopropanol. The genomic DNA was dissolved in nuclease-free water (2 μg/μl) and used as the template for amplification of the gRNA.

For sequencing of all gRNAs in the genome-wide library, the region containing the gRNA was amplified by a two-step PCR using Phanta Max Super-Fidelity DNA Polymerase (Vazyme, P505). In the first step, five 50-μl PCR reactions (each containing 40 μg of genomic DNA template) were performed with the forward P5-stagger primers mix (P5-stagger_0 to P5-stagger_8) and the reverse P7-stagger primers mix (P7-stagger_0 to P7-stagger_8). The staggered regions of different lengths in primers were added to maintain sequence diversity across the flow-cell. The thermal cycling parameters were 95 °C for 3 min, 16 cycles of 94 °C for 20 s, 55 °C for 15 s and 72 °C for 90 s, and the final extension, 72 °C for 10 min. Products of the first-step PCR were pooled together and used as the template for the second-step PCR. Six 50 μl reactions (each containing 1 μl of first-step PCR product) were performed with the forward primer P5-F and reverse primer P7-index. The thermal cycling parameters were 95 °C for 3 min, 20 cycles of 94 °C for 20 s, 55 °C for 15 s and 72 °C for 30 s, and the final extension, 72 °C for 5 min. The amplicons from the second-step PCR reactions were pooled and extracted on an agarose gel. The 150-bp paired-end sequencing was done with a HiSeq X Ten (Illumina). gRNA sequences were extracted and analysed with the CaRpools package (https://github.com/boutroslab/carpools). MAGeCK tool in this package was used for gene enrichment analysis (Fig. 3B).

### Generation of knockout or overexpression cell lines

To generate *SLFN11* knockout cell lines, HEK293, 293A and DU145 cells were transfected with px459 containing gRNA-1 (TATATCGCAAATATCCTGGT) or gRNA-2 (GTGAAGGTTATCGGGCGTTG) and then selected with puromycin (1 μg/ml) 48 hours post-transfection. Another 48 hours later, the cells were plated in a 96-well plate by serial dilutions. Single clones were screened by immunoblotting with SLFN11 antibody, and candidate knockout clones were verified by sequencing the PCR fragment from gRNA-targeted genomic regions. To generate *Slfn9* or *Tlr9* knockout B16-F10 cells, the Cas9 protein (IDT, #1081061) was incubated with synthetic targeted crRNA (IDT) (*Slfn9*: CGTTTGCGCCTCTACATGAG; *Tlr9*: CGCGTTCTCTTCATGGAC) and common tracrRNA (IDT) for 10 min and then electroporated into B16-F10 cells using the Neon Electroporation System (Thermo Fisher) following the manufacturer’s instructions. The cells were plated in a 96-well plate by serial dilutions. Single clones were screened by sequencing the PCR fragment from gRNA-targeted genomic regions.

Lentiviruses for overexpression Trex1 were produced in HEK293T cells as previously described. The HEK293 cells were infected with these viruses and then grown in a medium containing 1 μg/ml of puromycin to select for cells with stable expression.

### Subcellular fractionation

Nuclear and cytoplasmic extracts were prepared using the NE-PER Nuclear and Cytoplasmic Extraction Reagents Kit (Thermo Scientific, 78833) or Minute Cytoplasmic & Nuclear Extraction Kit for Cells (Invent, SC-003) at the appropriate time after transfection.

To dissect the cells into cytosolic, perinuclear and nuclear fractions (*34, 47*), cells (∼5×10^5^) from one well of 6-well plate were resuspended in the 200 μl of the hypotonic buffer RSB (10 mM NaCl, 10 mM Tris-HCl, pH 7.4, 1.5 mM MgCl2, with proteinase inhibitors) and incubated for 10 min on ice. The plasma membranes were further disrupted with 50 strokes of a glass Dounce homogenizer (loose pestle). After centrifugation at 1,000 × g for 5 min, the supernatant was further centrifuged at 10,000 × g for 10 min to obtain the cytosolic fraction. The nuclear plus perinuclear pellet was sequentially and gently washed with RSB and centrifuged at 1,000 × g for 5 min. After discarding the supernatant, the nuclear plus perinuclear pellet was resuspended in 200 μl of RSB containing 0.5% sodium deoxycholate and 1% Tween 40. The suspension was vortexed for 10 s and centrifuged at 1,600 × g for 5 min, and the supernatant was further centrifuged at 10,000 × g for 5 min to obtain the perinuclear fraction. The pellet consisting of the nuclear fraction was dissolved in 200 μl of nuclear extraction reagent (NER) (78835, Thermo Scientific) or in 200 μl of nuclear extraction buffer (SC-003-N, Invent). All steps were performed on ice or 4 °C.

For biotin-ODN pulldown and native ODN PAGE assay, cytosolic, perinuclear and nuclear fractions were extracted from 5-8×10^6^ cells cultured on 10-cm dishes following the steps described above using 1 ml of the appropriate buffer for each fraction.

### Luminous ODIP and biotin-ODN pulldown assays

Approximately 3 × 10^6^ cells were transfected with 5 μg of biotinylated ODNs complexed with Lipofectamine 3000. After 6-12 hours of transfection, the cells were washed twice with PBS.

For the ODIP assay, 3 × 10^6^ cells were lysed in 500 μl of IP lysis buffer (Beyotime, P0013) containing protease inhibitor. After centrifugation, the cell lysates were incubated with the appropriate IgG and Protein G magnetic beads (Thermo Scientific, 88847) for 2 hours at 4 °C for pre-clearing. The beads were then discarded, and 1.5 μg of SLFN11 (D-2) antibody was added to the cell lysates, which were incubated overnight at 4°C. Next, 15 μl of protein G magnetic beads was added to the cell lysates and incubated for 2.5 hours. After washing with PBS for three times, the beads were resuspended in 70 μl of stable peroxide solution (Thermo Scientific, 89880) and transferred to a well in a 96-well plate. The chemiluminescent signal was recorded using Luminescent Imaging Workstation (Tanon) after adding 70 μl of luminol/enhancer solution. The beads were then recovered in a new tube, and the proteins were boiled off the beads and detected by immunoblotting.

For biotin-ODN pulldown in whole cell lysate, 3 × 10^6^ cells were lysed in 500 μl of Buffer L (50 mM Tris-HCl (pH 8.0), 150 mM KCl, 1 mM DTT, 0.5% Triton X-100, 10% glycerol) containing protease inhibitor. After centrifugation, the cell lysates were incubated with 15 μl of SA beads (Invitrogen, 65601; Merck Millipore, 69203) for 6 hours at 4 °C. The supernatants were discarded, and the beads were washed with Buffer L once and with washing buffer (20 mM Tris-HCl (pH 7.5), 100 mM KCl, and 2 mM EDTA) twice. The protein/ODN complexes were boiled off the beads and detected by immunoblotting as described above. To perform biotin-ODN pulldown assays from cytosolic, perinuclear and nuclear fractionations, 25 μl of SA beads was added to each 1-ml fraction, and the steps described above were followed.

For biotin-ODN pulldown in cytosolic, perinuclear and nuclear fractions, 5 × 10^6^ cells were transfected with 9 μg of biotinylated ODNs. After 8 hours of transfection, the cells were washed twice with PBS and dissected into three fractions. The biotin-ODN pulldown assay followed the steps described above.

### Denaturing and native ODN PAGE

The cells were transfected with biotinylated ODNs. For denaturing ODN PAGE, the cells were lysed in SNET buffer containing proteinase K and incubated at 55 °C overnight. The ODNs in proteinase K-digested lysates were denatured for 5 min at 95 °C and then snap-cooled on ice. Samples were loaded onto an 8% or 12% polyacrylamide gel containing 8 M urea. The gels were run at 12.5 V/cm at room temperature. For native ODN PAGE, the cells were dissected into the indicated subcellular fractions as previously described. Samples were separated by electrophoresis through native polyacrylamide gels (6%) in 0.5×TBE buffer at 12.5 V/cm and 4 °C. For the detection of biotinylated ODNs, ODNs were electrophoretically transferred to a nylon membrane at 250 mA for 45 min. After UV-light crosslinking, the biotinylated ODNs were detected with the Chemiluminescent Nucleic Acid Detection Module Kit (Thermo Fisher, 89880) according to the manufacturer’s user guide.

### Purification of recombinant proteins

To express GST-tagged C-terminal SLFN11 in bacteria, *E. coli* BL21 (DE3) cells containing pGEX-SLFN11 (residues 349-901) plasmids were grown in Luria-Bertani medium supplemented with ampicillin. Protein expression was induced overnight at 18 or 37 °C with 0.4 mM isopropyl-B-D-thiogalactopyranoside (IPTG). As SLFN11 was in an aggregated state in inclusion bodies, the inclusion bodies were purified from sonicated cell lysates and solubilized with extraction buffer (20 mM NaHCO_3_, 50 mM NaCl, 0.5 mM EDTA, 0.5% sodium lauroyl sarcosine (SKL), 5 mM DTT). The solubilized SLFN11 was refolded with redox buffer (1 mM oxidized glutathione, 2 mM reduced glutathione and 0.2% PEG 4000). To express His-tagged C-terminal SLFN11 in insect cells, the Bac-to-Bac baculovirus expression system (Thermo Fisher) was used following the manufacture’s instructions. Sf9 cells (1×10^7^) were transfected with 10 µg of pFastbac1 plasmid containing the SLFN11 (residues 349-901) coding frame to produce baculovirus. One litre of Sf9 cells was infected with 10 ml of the P0 baculovirus and cultured at 28 °C for 3 days for large-scale expression. His-tagged SLFN11 (residues 349-901) was purified by Ni-NTA beads (Qiagen). All purified GST- or His-tagged proteins were dialyzed against a buffer containing 50 mM Tris-HCl and 150 nM NaCl and validated by Coomassie blue-stained SDSLJPAGE.

### Binding affinity measurements by biolayer interferometry (BLI)

All BLI measurements were performed on a ForteBio’s Octet RED96 system using BLI buffer (50 mM Tris-HCl (pH 7.5), 150 mM KCl, 2 mM MgCl2, 1 mM DTT, 6% glycerol, 0.02% Tween 20). To measure the direct binding between GST-c-SLFN11 (residues 349-901) and CpG ODNs, GST-c-SLFN11 was immobilized onto Anti-GST Biosensor tips (Sartorius) in BLI buffer by dipping the biosensors into wells containing GST-c-SLFN11 for 300 s. After obtaining the baseline reading, the biosensors were next transferred to wells containing the indicated biotinylated ODNs to monitor the association of ODNs with immobilized GST-c-SLFN11 for 600 s. Finally, the biosensors were transferred to wells containing BLI buffer to monitor the dissociation of ODNs from GST-c-SLFN11 protein for 600 s.

To determine the kinetics of the interactions of ODN-H with His-c-SLFN11, biotinylated ODN-H was immobilized onto SA biosensor tips (Sartorius), and His-c-SLFN11 was added into wells for the association step. The dissociation constant (Kd) values were obtained by fitting the binding data globally to a 1:1 binding model using Data Analysis software 11.1 from ForteBio.

### EMSA and fluorescence ODN GST pulldown assays

For the EMSA assay (fig. S6E), 0.2 μg of purified His-c-SLFN11 (residues 349-901) protein was incubated with 20 pmol of the indicated FAM-labelled CpG ODNs in binding buffer (5LJmM Tris pHLJ7.5, 25 mM KCl, 0.5 mM dithiothreitol (DTT), 5 mM MgCl2, 0.05% NP-40, 10 mM EDTA, 2.5% glycerol, 0.05 mg/ml poly (deoxyinosinic-deoxycytidylic) (dI-dC)) in a 20 μl reaction system for 30 min. The samples were then loaded onto a 5% native polyacrylamide gel that had been pre-run in 0.5× TBE (Tris-borate-EDTA) buffer for 60LJmin, and ran at constant voltage of 100 V in 0.5× TBE at 4 °C. The gel was then scanned using Typhoon FLA 9500 (GE) in fluorescence detection mode (LD laser).

For the fluorescence ODN GST pulldown assay (fig. S6F), an assay based on ODIP, GST-c-SLFN11 (0.5 μg) was incubated with FAM-labelled CpG ODNs (50 pmol) in 50 μl of BLI buffer for 1 hours at RT. Glutathione agarose beads (Pierce) (15 μl) were washed twice with BLI buffer and then added to the GST-c-SLFN11 and ODN mixtures for another 1 hours of incubation. After discarding the supernatant, the beads were washed twice with BLI buffer and transferred to a 96-well plate with clear bottom. The 96-well plate was scanned as described for the EMSA gel, and the fluorescence values of the wells were also recorded by Cytation 5 (BioTek).

### Immunofluorescence assay

HEK293 or DU145 cells were seeded on coverslips in 24-well plates and transfected with the indicated biotinylated ODNs. After 8-12 hours of transfection, the cells were washed with PBS and fixed with 4% paraformaldehyde (PFA) in PBS for 15 min at room temperature or overnight at 4 °C. The cells were permeabilized for 10 min with PBS containing 0.2% Triton X-100. Biotinylated ODNs were stained using the Tyramide SuperBoost kit (Thermo Fisher, B40935) according to the manufacturer’s introduction. In detail, the endogenous peroxidase activity of the cells was quenched by incubation with a 3% hydrogen peroxide solution for 1 hours, followed by washing steps with PBS. The cells were then blocked for 1 hours with blocking buffer and stained with HRP-conjugated SA for 1 hours, which reacted with Alexa Fluor tyramide to provide the fluorescence signal. After ODN staining, the cells were incubated overnight with primary antibodies/5% BSA/PBS at a 1:300 dilution for SLFN11 (D-2) (*29*). After washing with PBST, the cells were incubated with anti-mouse secondary antibodies and stained with Hoechst 33342. When ODN detection was not necessary, the cells were incubated with the primary antibody, after the blocking step. For intracellular ssDNA staining, HEK293 cells were treated with 4-mM HU for 8 hours, and then stained with anti-ssDNA antibody following the steps described above. Images were captured with an Olympus FV1000 or a Zeiss LSM 780 confocal microscope. All image data shown are representative of 5-10 randomly selected fields. The number of cells with SLFN11 cytosolic translocation was recorded from 150 HEK293 cells or 100 DU145 cells and the repeated times were indicated (fig. S8).

### Nanoparticle formation

LNPs containing CGT ODNs, heat-denatured AAV2 gDNA or firefly luciferase mRNA (APExBIO, R1018) were prepared using the ethanol dilution method (*48*). In brief, ALC-0315 ([(4-hydroxybutyl) azanediyl] di (hexane-6,1-diyl) bis (2-hexyldecanoate)), 1,2-dioleoyl-sn-glycero-3-phosphocholine (DOPC), cholesterol, and 1,2-dimyristoyl-rac-glycero-3-methoxypolyethylene glycol-2000 (DMG-PEG 2000) were dissolved in ethanol at a molar ratio of 50: 18.5: 30: 1.5. The indicated type of nucleic acids was prepared in 25 mM citrate sodium buffer at pH 4.0. The two solutions were rapidly mixed at an aqueous to ethanol ratio of 3:1 (vol./vol.) to achieve a final weight ratio of 40/1 (total lipids/nucleic acids), and then incubated for 15 min at room temperature. The ethanol and unloaded nucleic acids in the mixtures were removed using Amicon Ultra centrifugal filters (10- or 30-kD MWCO, Merck) for two rounds with 5- and 15-times volumes of 1× PBS, respectively. The theoretical nucleic acid-to-lipid ratios for all formulations were maintained at an N/P charge ratio (the ratio of the charge on the cationic lipid, assuming that it was in the positively charged protonated form, to the negative charge on the nucleic acids) of 3. Each prepared LNP sample was adjusted to ∼0.6 mg/mL nucleic acids with 1× PBS for storage at 4 °C.

### Animal and in vivo CpG ODN delivery

All animal protocols were approved by the Animal Care and Use Committee at the Institute of Basic Medical Sciences, Chinese Academy of Medical Sciences, and Peking Union Medical College. C57BL/6 WT mice were purchased from the Vital River Laboratory Animal Technology Co. and *Tlr9^-/-^* mice were a gift from Zhigang Tian (China Science & Technology University) and originally generated by Shizuo Akira (*26*). *Slfn9* knockout mice were generated by microinjection of in vitro-translated Cas9 mRNA and a pair of gRNAs into C57BL/6 zygotes. The gRNA sequences used are shown (fig. S9A). Founders were validated by DNA sequencing. A founder with a 31-bp deletion in the coding region was chosen to intercross with the WT mice to produce F1 *Slfn9^+/-^*mice. Every generation of *Slfn9^+/-^* mice was intercrossed with WT mice until the F3 generation to eliminate potential off-targets. *Slfn9^-/-^*mice were obtained by intercrossing F3 *Slfn9^+/-^* mice.

Neonatal mouse experiments were conducted using one-to-two-day-old pups. To assess the tissue distribution of LNPs, 50 μl of PBS containing 2 μg of Fluc mRNA encapsulated in LNPs was injected through the temporal vein. Pups were injected intraperitoneally with D luciferin (150 mg/kg) at 6 hours after LNP injection and imaged using an IVIS Lumina system (Perkin Elmer). To evaluate the in vivo response to CGT ODNs, pups were injected with 50 μl of PBS containing 6 μg of HIV6441-LNPs through the temporal vein. Liver tissues were collected 6 hours post-injection for qPCR and immunoblotting assays.

For juvenile (3-4 weeks old) and adult (10-12 weeks old) mice, 3 mg/kg HIV6441-LNPs or AAV2 gDNA-LNPs were injected through the tail vein. Liver tissues were collected 6 hours post-injection for qPCR assays and 72 hours post-injection for histological assays and ALT and AST measurement. Shock assays were conducted on juvenile and adult mice injected with 3 mg/kg AAV2 gDNA-LNPs and 5 mg/kg HIV6441-LNPs, respectively, and they were monitored four times daily for a total of four days.

### Antitumor monotherapy

B16-F10 cells (2×10^5^) in 100 μl of PBS were implanted into right flank of 6-week-old C57BL/6 female mice. HIV6441 was incubated with Entranster-in vivo (Engreen) following the manufacturer’s protocol. The solid tumors that formed were injected with 10 μg of HIV6441 on days 4, 6, 8 and 10. Tumors were examined under a fiber optic illuminator and its long (L) and short (W) diameters were measured every 3-4 days with a caliper. Tumor volume was determined using the volume formula for an ellipsoid. Mice were sacrificed by CO_2_ inhalation when the tumor volume reached 2,000 mm (*49*).

### Statistical analyses

Experiments were independently repeated two to four times. For in vivo experiments, mice were randomly assigned to groups and mixed among cages. Student’s t tests, log-rank tests and one-way and two-way ANOVA with Bonferroni correction for multiple tests were performed using Prism software (Graphpad version 9.5.1). The statistical methods used for each analysis are specified in the figure legends, and *P* < 0.05 was considered statistically significant.

